# REC114 partner ANKRD31 controls number, timing and location of meiotic DNA breaks

**DOI:** 10.1101/425322

**Authors:** Michiel Boekhout, Mehmet E. Karasu, Juncheng Wang, Laurent Acquaviva, Florencia Pratto, Kevin Brick, Diana Y. Eng, R. Daniel Camerini-Otero, Dinshaw J. Patel, Scott Keeney

**Affiliations:** Molecular Biology Program, Memorial Sloan Kettering Cancer Center, New York, NY, USA; Louis V. Gerstner, Jr. Graduate School of Biomedical Sciences, Memorial Sloan Kettering Cancer Center, New York, NY, USA; Structural Biology Program, Memorial Sloan Kettering Cancer Center, New York, NY, USA; Genetics & Biochemistry Branch, NIDDK, National Institutes of Health, Bethesda, MD, USA; Howard Hughes Medical Institute, Memorial Sloan Kettering Cancer Center, New York, NY, USA

## Abstract

Double-strand breaks (DSBs) initiate the homologous recombination that is crucial for meiotic chromosome pairing and segregation. Here we unveil mouse ANKRD31 as a lynchpin governing multiple aspects of DSB formation. Spermatocytes lacking ANKRD31 have altered DSB locations and fail to target DSBs to sex chromosomes’ pseudoautosomal regions (PAR). They also have delayed/fewer recombination sites but, paradoxically, more DSBs, suggesting DSB dysregulation. Unrepaired DSBs and pairing failures—stochastic on autosomes, nearly absolute on X and Y—cause meiotic arrest and sterility in males. *Ankrd31*-deficient females have reduced oocyte reserves. A crystal structure defines direct ANKRD31– REC114 molecular contacts and reveals a surprising pleckstrin homology domain in REC114. *In vivo*, ANKRD31 stabilizes REC114 association with the PAR and elsewhere. Our findings inform a model that ANKRD31 is a scaffold anchoring REC114 and other factors to specific genomic locations, promoting efficient and timely DSB formation but possibly also suppressing formation of clustered DSBs.

## Introduction

Meiotic recombination initiates with developmentally programmed DSBs (de Massy, 2013; Lam and Keeney, 2014). The potentially lethal nature of these DSBs makes it important for cells to control their timing, number, and location (Keeney et al., 2014), but how DSBs form and how they are controlled remain poorly understood, particularly in mammals.

Meiotic DSBs are made by the topoisomerase-like SPO11 protein in conjunction with a suite of accessory proteins (“DSB proteins”), most but not all of which are evolutionarily conserved (Keeney, 2008; Lam and Keeney, 2014; Robert et al., 2016). Spo11 has nine essential partners in budding yeast, where DSB proteins are best characterized, including the meiosis-specific Rec114, Mei4, and Mer2. In mice, DSB formation similarly requires REC114, MEI4, and IHO1 (a Mer2 ortholog) (Kumar et al., 2010; Stanzione et al., 2016; Tesse et al., 2017; Kumar et al., 2018). We have a broad-strokes picture of the network of interactions connecting these proteins (e.g., Arora et al., 2004; Li et al., 2006a; Maleki et al., 2007; Miyoshi et al., 2012), but mechanistic detail is lacking. How these proteins localize to chromosomes is also not well defined. Moreover, while many of the DSB proteins are nearly universal in eukaryotes, phylogenetically restricted examples such as vertebrate MEI1 (Libby et al., 2002) raise the possibility that the catalog of relevant proteins may not be complete.

DSB number and timing are controlled by intersecting negative feedback circuits (Cooper et al., 2014; Keeney et al., 2014). In one circuit, activation of the DNA damage-response kinase ATM inhibits further DSB formation locally, preventing multiple breaks nearby on the same chromatid and on the sister chromatid (Lange et al., 2011; Zhang et al., 2011; Carballo et al., 2013; Anderson et al., 2015; Garcia et al., 2015). In another circuit, chromosomes that successfully engage their homologs stop making DSBs, likely via displacement of pro-DSB factors (Wojtasz et al., 2009; Carballo et al., 2013; Kauppi et al., 2013; Thacker et al., 2014). These and other regulatory circuits collaborate to ensure that sufficient DSBs are made to support homologous chromosome pairing and recombination without a deleterious excess of DNA damage (Cooper et al., 2014; Keeney et al., 2014).

DSB locations are also controlled, making recombination distributions highly nonrandom (Baudat et al., 2013; Lam and Keeney, 2014). In most mammals, a critical component of this control is PRDM9, which contains a sequence-specific DNA binding domain and a PR/SET histone methyltransferase domain that methylates histone H3 on lysines 4 and 36 (Baudat et al., 2010; Myers et al., 2010; Wu et al., 2013; Powers et al., 2016; Diagouraga et al., 2018). PRDM9 has thousands of recognition sites across the genome. Binding to these sites results in methylation of nearby nucleosomes and subsequent targeting of SPO11 to form DSB hotspots. In the absence of PRDM9, SPO11 instead preferentially targets “default” sites, often where PRDM9-independent H3K4 trimethylation occurs such as promoters and CpG islands (Brick et al., 2012). There is also a handful of sites targeted independently of PRDM9 in normal meiosis (Brick et al., 2012). Perhaps the most prominent of these is the pseudoautosomal region (PAR), the small segment where the X and Y chromosomes share homology and must recombine to ensure sex chromosome segregation in male meiosis (Rouyer et al., 1986; Soriano et al., 1987). Although the central roles of PRDM9 DNA binding and histone methylation are clear, the downstream factors that connect these hotspot designation events to SPO11 activity remain to be identified (Grey et al., 2018). The mechanisms of PRDM9-independent SPO11 targeting are likewise poorly understood.

Here we relate how a search for mammalian meiotic recombination proteins identified ANKRD31 (<u>ank</u>yrin <u>r</u>epeat <u>d</u>omain-containing 31) as a previously unknown but critical component of the DSB-targeting and control machinery. ANKRD31 is essential for male meiosis because it regulates the timing, number, and location of DSBs globally and is crucial for DSB formation specifically in the PAR. Our findings show that ANKRD31 stabilizes the association of pro-DSB factors with chromosomes by direct interaction with REC114, details of which are revealed by atomic resolution structural analysis.

## Results

### ANKRD31 is a meiosis-specific, chromatin-associated, REC114-interacting protein

To identify proteins involved in meiotic DSB formation, we used mouse REC114 and MEI4 as yeast two-hybrid baits to screen a cDNA library from leptotene/zygotene spermatocytes. REC114 was recovered when MEI4 was the bait, consistent with known interactions in mouse (Kumar et al., 2010) and yeasts (Arora et al., 2004; Miyoshi et al., 2012). When REC114 was the bait, the screen yielded two clones encoding portions of the previously uncharacterized protein ANKRD31 (**Figures 1A and 1B**), among other hits (see Methods).

**Figure 1.**
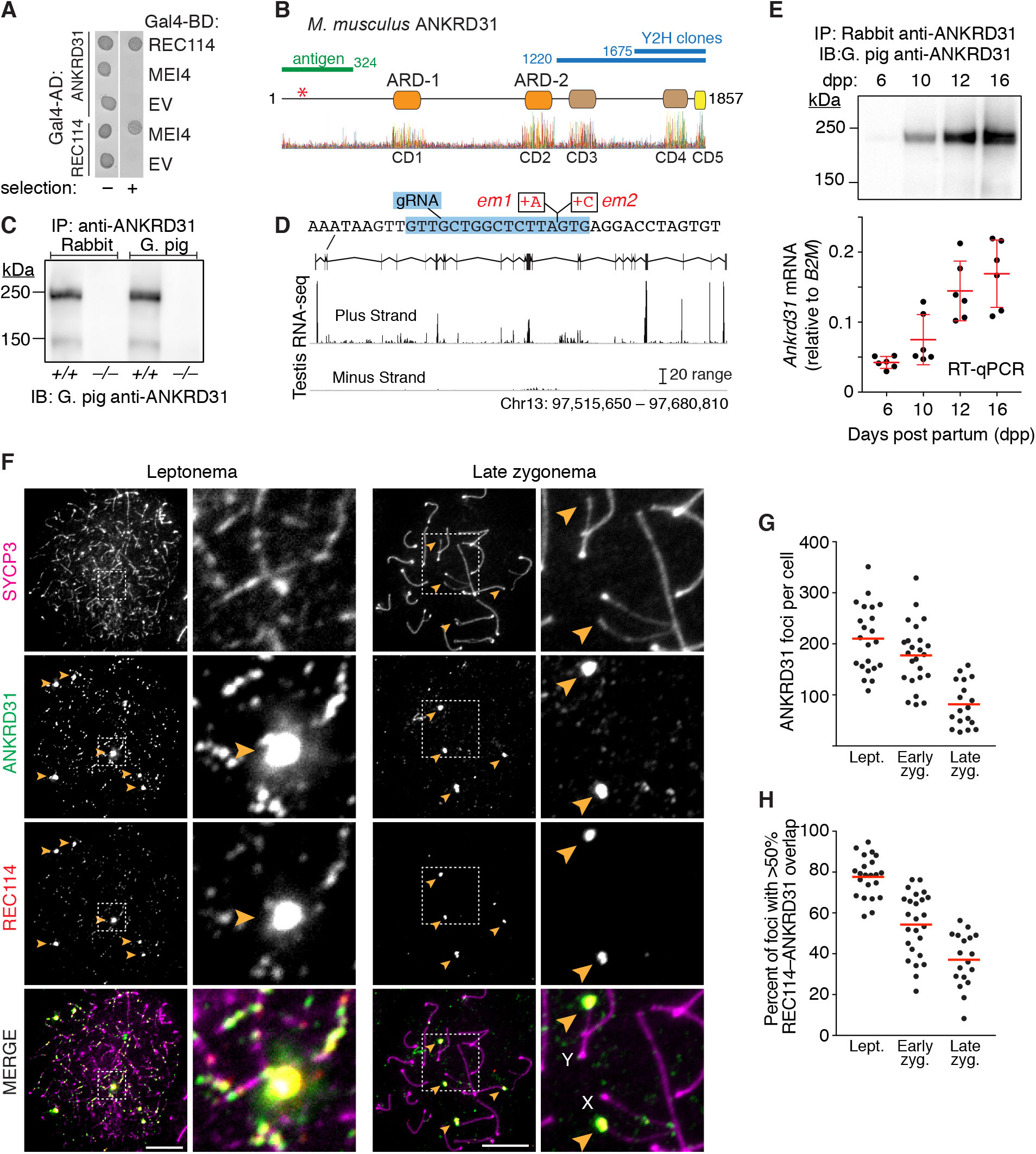
ANKRD31 is a meiosis-specific REC114-interacting protein. (A) Yeast two-hybrid assay showing REC114 interactions with ANKRD31 and MEI4. Growth of cells expressing the indicated Gal4 activating domain (AD) and binding domain (BD) fusions is shown. EV, empty vector. “Selection” indicates presence of aureobasidin to detect activation of the two-hybrid reporter. (B) Domain structure of mouse ANKRD31 (above) and conservation plot for vertebrate homologs (below, (Wheeler et al., 2014)). Positions of frameshift mutation (asterisk), yeast two-hybrid (Y2H) clones, and antigen used for antibody production are indicated. Conserved domains (CD1–CD5) are indicated. (C) Immunoprecipitation (IP) and immunoblotting (IB) of ANKRD31 from whole-testis extracts using antisera raised in the indicated species. (D) Ankrd31 exon map with browser shot showing ENCODE long RNA-sequencing reads from adult testis. Sequence context for the CRISPR-Cas9 guide RNA (gRNA in blue) and frameshift mutations in exon 3 (em1 and em2) is shown. (E) Expression time course of ANKRD31 protein (top, IP/IB) and RNA (below, reverse-transcription quantitative PCR) in testes of juvenile mice. RT-qPCR was normalized to expression of the B2M gene. Red lines, mean ± s.d. (F) ANKRD31 localizes to chromatin. Spread spermatocyte chromosomes were stained for SYCP3, ANKRD31 (guinea pig antiserum), and REC114. Arrowheads indicate examples of ANKRD31 blobs. The zoomed images show ANKRD31 colocalization with REC114, both in small foci and in larger blobs. Scale bars are 10 μm. (G) ANKRD31 focus counts at different meiotic stages (n = 2 mice). (H) Colocalization of ANKRD31 with REC114. See also Figure S1

To prioritize these candidates, we evaluated sequence conservation. Meiotic factors are often rapidly evolving (Swanson and Vacquier, 2002; Keeney, 2008). For example, the meiotic DNA strand exchange protein DMC1 and the meiotic cohesin subunits REC8, RAD21L, and SMC1β have diverged more than their paralogs that function in all cells (RAD51, RAD21, and SMC1α) (**Figure S1A**). We reasoned that proteins displaying this signature might be directly involved in meiotic processes. We therefore focused on ANKRD31 because it has relatively high rates of sequence divergence in mammals (**Figure S1B**).

*Ankrd31* cDNA (5.9 kb) from testis matched NCBI reference RNA XM_006517797.1, comprising 26 exons and encoding a predicted protein of 206 kDa (**Figure 1B**). ANKRD31 has two ankyrin-repeat domains (ARDs) with three ankyrin repeats apiece. Each repeat is a ~33-amino-acid alpha-helical motif and ARDs are often involved in protein-protein interactions (Li et al., 2006b). We identified likely orthologs across the vertebrate subphylum including cartilaginous fish (e.g., ghost shark *Callorhinchus milii*), but not in more distant taxa. Alignments of vertebrate orthologs identified conserved domains (CDs) overlapping each ARD (the first appears to be absent in fish species) plus three CDs in the C-terminal third of the protein (**Figure 1B**). CD5 at the C-terminus interacts directly with REC114 (detailed below); functions of the others remain to be determined.

Polyclonal antisera raised in rabbit and guinea pig recognized a protein migrating near the expected size from extracts of adult testes, and this protein was absent from homozygotes for an *Ankrd31* frameshift allele described below, confirming specificity (**Figure 1C**). *Ankrd31* protein and mRNA were present in adult testis with little or no expression in somatic tissues (**Figures 1D, S1C and S1D**), and they accumulated during the first wave of meiosis in testes of juvenile mice (**Figure 1E**). *Ankrd31* expression thus appears to be largely meiosis-specific.

ANKRD31 is chromatin-associated, based on immunostaining of spread spermatocyte chromosomes (**Figure 1F**). Early in prophase I, contemporaneous with recombination initiation, aligned sister chromatids begin to form an SYCP3-containing protein axis from which chromatin emanates in loops (leptonema). The axes elongate and homologous chromosomes begin to align and form stretches of synaptonemal complex (SC), a tripartite structure comprising the homologous axial elements plus the central region connecting them (zygonema). The SC elongates until it juxtaposes each homolog pair along their lengths (pachynema), then disassembles as cells prepare to exit prophase (diplonema).

At leptonema, ANKRD31 formed numerous foci mostly decorating chromosome axes (average of 210 per cell; **Figure 1F**). Focus numbers then decreased progressively through zygonema, particularly in synapsed regions (**Figures 1F, 1G and S1E**). Little or no background staining was seen in the *Ankrd31* mutant, again indicating antibody specificity (**Figure S1F**). Several heavily-stained ANKRD31 blobs were also detected (arrowheads in **Figure 1F**), which correspond to the PARs of the X and Y chromosomes (see late zygotene cell in **Figure 1F**) and other regions that have PAR-like sequences containing arrays of the mo-2 minisatellite (Acquaviva, Jasin & Keeney, unpublished). ANKRD31 colocalized with REC114, both in small foci and in the mo-2-associated blobs (**Figures 1F, 1H, and S1E**), supporting the idea that these proteins interact physically in vivo. ANKRD31 also colocalizes with MEI4, MEI1, and IHO1 (Acquaviva, Jasin & Keeney, unpublished).

### Ankrd31 mutation causes male sterility from spermatogenic arrest

To determine the function of *Ankrd31*, we used CRISPR-Cas9 to generate endonuclease-mediated (em) mutations in exon 3. Two alleles were recovered with single-nucleotide insertions at the same position (**Figure 1D**). Indistinguishable phenotypes were observed for either homozygote where tested, so for simplicity experiments with the *Ankrd31^em1^* allele (hereafter *Ankrd31^−^*) are shown unless indicated otherwise. *Ankrd31^−/−^* testes lacked detectable signal in IP/westerns (**Figure 1C**), indicating these are null or severe loss-of-function alleles. Distinct mutations generated by Tóth and colleagues gave similar phenotypes (Papanikos, Tóth et al., personal communication), reinforcing that these alleles abolish ANKRD31 function.

*Ankrd31^+/−^* heterozygotes showed normal fertility and Mendelian transmission of the mutations (**Figure S2A**). Histopathological analysis turned up no obvious somatic defects in homozygous mutants (see Methods), suggesting that ANKRD31 is dispensable in non-meiotic cells. However, *Ankrd31^−/−^* males were sterile: none of the 4 animals tested sired offspring when bred with wild-type females for 16 weeks, and testes were 37% of the size in littermate controls (**Figures 2A and S2B**).

**Figure 2.**
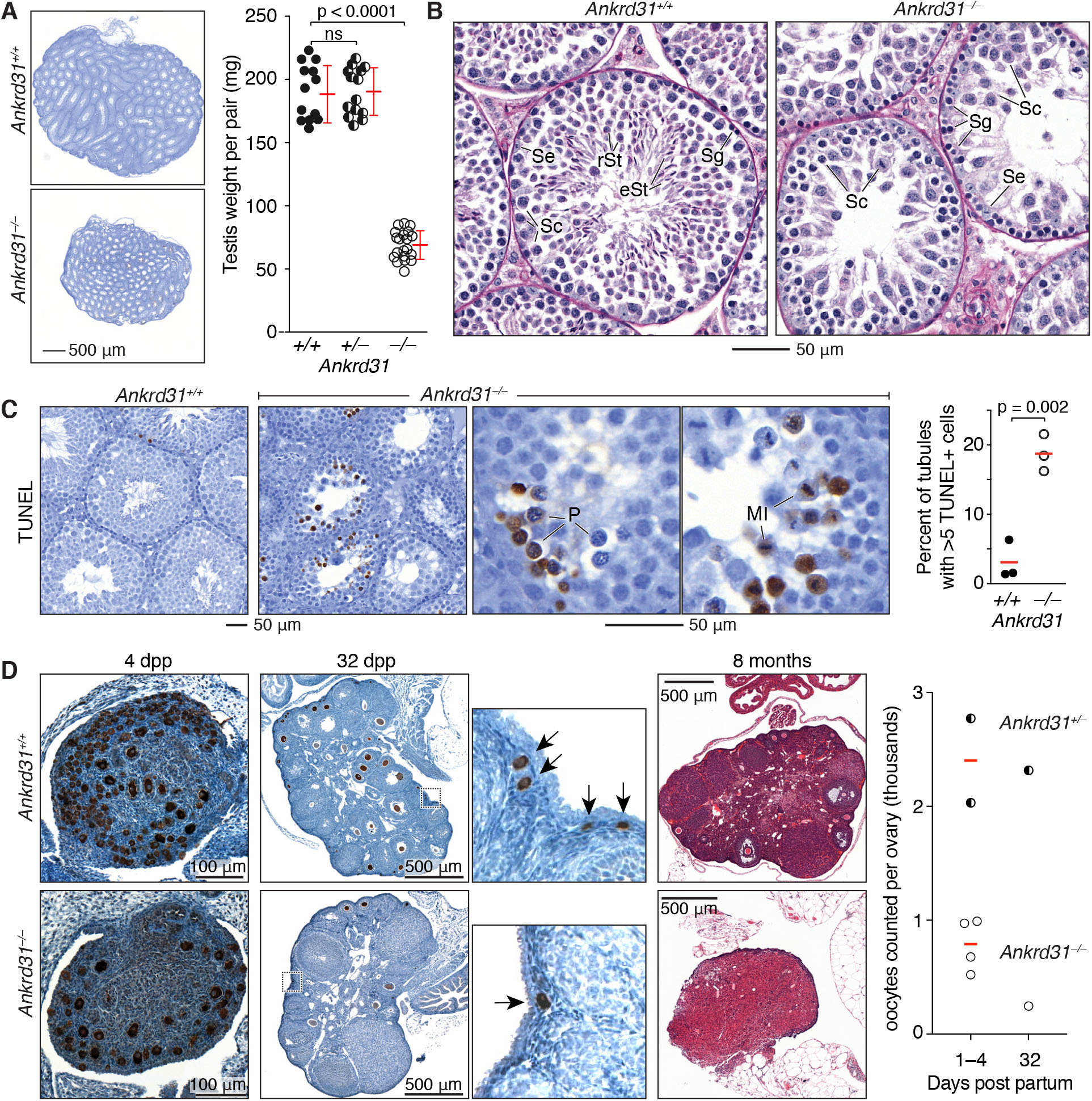
*Ankrd31* in male and female fertility. (A) Reduced testis size in *Ankrd31^−/−^* mutants. Representative sections from adult testes are shown at left (4 mos old); quantification of adult testis weights is at right (red lines, mean ± s.d.). (B) Defective spermatogenesis in *Ankrd31^−/−^* mutants. Representative Bouin’s-fixed and periodic acid Schiff (PAS)-stained seminiferous tubule sections from adult testes are shown (5 mos old). Se, Sertoli cells; Sg, spermatogonia; Sc, spermatocytes; rSt, round spermatids; eSt, elongated spermatids. (C) Increased apoptosis in *Ankrd31^−/−^* spermatocytes. Lower and higher magnification images (left and right panels, respectively) are shown of adult testis sections stained with TUNEL and counterstained with hematoxylin (4 mos old). Each point on the graph is the measurement from one animal; red line indicates mean. (D) Reduced oocyte reserve and premature ovarian failure in *Ankrd31*-deficient females. Representative ovary sections at the indicated ages are shown. Samples at 4 and 32 dpp were immunostained for MVH to mark oocytes (brown stain); insets for 32 dpp ovaries highlight primordial follicles (arrows) near the cortex. The samples at 8 mos were Bouin’s fixed and PAS-stained. The graph at right shows oocyte counts summed across every third serial section. Red lines indicate means. In panels A and B, the results of two-tailed t tests are indicated (ns, not significant (p > 0.05). See also Figure S2.

Male mice unable to initiate or complete meiotic recombination display sterility and hypogonadism because of apoptotic elimination of spermatocytes (de Rooij and de Boer, 2003). To determine when gametogenesis fails in *Ankrd31* mutants, we examined testis sections. Control animals had the full array of spermatogenic cells including spermatocytes and round and elongated spermatids, as expected (**Figure 2B**). In contrast, *Ankrd31^−/−^* tubules contained spermatogonia and primary spermatocytes but were largely if not completely devoid of postmeiotic cells (**Figure 2B**). TUNEL staining detected an elevated frequency of apoptosis (**Figures 2A and 2C**). In a subset of apoptotic tubules, dying cells were pachytene spermatocytes (**Figure 2C**), similar to mutants lacking or unable to repair DSBs (e.g., *Spo11^−/−^* or *Dmc1^−/−^*) (Barchi et al., 2005). However, pachytene arrest was only partially penetrant, as subsets of tubules instead contained apoptotic cells at metaphase I (**Figure 2C**).

### Reduced oocyte reserve and premature ovarian failure in Ankrd31^−/−^ females

Young adult *Ankrd31^−/−^* females were fertile (**Figure S2A**), but the partially penetrant pachytene arrest in males led us to consider that similar stochastic prophase failure might occur in females. Oocytes initiate meiosis during fetal development and finish pachynema around birth, then arrest and form resting follicles. A large fraction of wild-type oocytes fail to make follicles, but meiotic prophase defects cause even greater oocyte loss (Hunter, 2017).

If ANKRD31 contributes to female meiosis, we therefore predicted more oocyte culling and fewer follicles. To test this, we examined ovaries at various ages (**Figures 2D and S2C**). As expected, wild-type animals had abundant oocytes or follicles at 1 and 4 days post partum (dpp) and a mix of resting (primordial) and growing follicles at 32 dpp. In contrast, *Ankrd31^−/−^* females had greatly reduced oocyte numbers at all ages examined. Growing follicles were observed at 32 dpp, so ANKRD31 is dispensable for follicle development per se, but the mutant had fewer primordial follicles indicating a smaller oocyte reserve (**Figure 2D** insets). Exhaustion of this reserve caused reduced ovary size by ~8 mos, suggesting premature ovarian failure (**Figure 2D**). This partially penetrant oocyte elimination shows that ANKRD31 is required for normal female fertility.

### ANKRD31 deficiency causes stochastic synapsis failures

To understand ANKRD31 function, we focused on the mutant phenotype in males. The unusual mixed arrest—partial in pachynema and more complete in metaphase I—suggested that the protein might contribute to multiple processes. We therefore addressed each arrest in detail. Pachytene apoptosis can be triggered by persistent DSBs or by failure to transcriptionally silence the sex chromosomes, both of which can also be associated with defects in homologous synapsis (Burgoyne et al., 2009; Royo et al., 2010; Pacheco et al., 2015).

We tracked synapsis by immunostaining spread spermatocyte chromosomes for SYCP3 and the SC central region component SYCP1 (de Vries et al., 2005). In *Ankrd31^−/−^* mutants, leptonema and zygonema appeared similar to wild type and most pachytene cells had apparently complete autosomal synapsis, thus ANKRD31 is dispensable for formation of axial elements and SC per se (**Figures 3A, 3B and S3A**). However, 14% of pachytene-like cells displayed a mix of fully synapsed autosomes plus asynaptic chromosomes and/or chromosome tangles, i.e., a mix of synapsed and unsynapsed axes with partner switches indicative of nonhomologous synapsis (**Figures 3A, 3B and 3C**). Thus some *Ankrd31^−/−^* cells appear to have homologous pairing defects. Synapsis defects of this sort can cause apoptosis (Turner et al., 2006; Kauppi et al., 2013), so stochastic SC failures may contribute to death of some mutant spermatocytes in pachynema. Other chromosome structure abnormalities were also observed, including inter- and intrachromosomal end-to-end fusions (**Figure S3B**).

**Figure 3.**
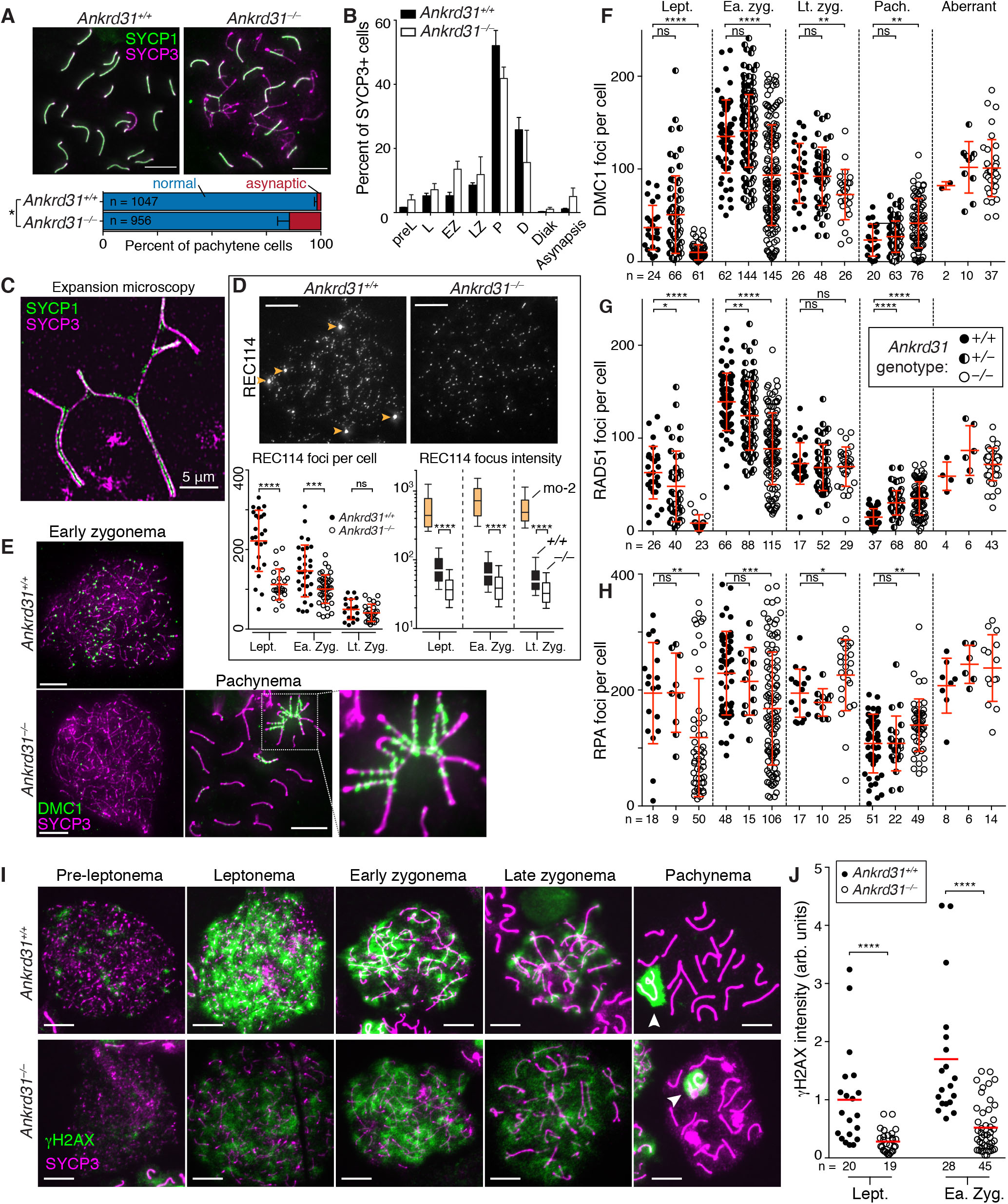
Dysregulated recombination in *Ankrd31^−/−^* spermatocytes. (A) Synapsis defects in a subset of pachytene cells. Above, representative chromosome spreads stained for SYCP3 and SYCP1; below, quantification of synapsis defects from two animals of each genotype (bars indicate mean and range, n is total number of pachytene cells counted). (B) Quantification of spermatocyte stages based on staining for SYCP3 and γH2AX (pre-leptonema, leptonema, early zygonema, late zygonema, pachynema, diplonema, diakinesis, respectively; pachytene-like cells with autosome asynapsis were tallied separately). Bars indicate mean and range of two animals per genotype. (C) Expansion microscopy showing SC partner switches in a pachytene-like *Ankrd31^−/−^* spermatocyte. (D) ANKRD31 is required for normal REC114 localization to chromosomes. Representative micrographs at matched exposures are shown above. Arrowheads indicate REC114 blobs, which are missing in the *Ankrd31^−/−^* mutant. Each point in the graph at lower left is the REC114 focus count for a cell of the indicated stage (red lines indicate mean ± s.d.). The box plot at lower right summarizes REC114 focus intensities from Experiment 1 in Figure S3C. Boxes indicate median, 25th and 75th percentiles; whiskers indicate 10th and 90th percentiles; outliers are not shown. Intensities of the blobs associated with mo-2 in wild type are summarized separately for comparison (tan boxes). Intensities of the smaller foci were lower in *Ankrd31^−/−^* (open boxes) than in wild type (black filled boxes) at all stages. (E–H) Altered numbers and timing of recombination foci. Representative images of DMC1 staining are shown in E; quantification of DMC1, RAD51, and RPA focus counts are shown in F, G, and H, respectively. “Aberrant” refers to pachytene-like cells with unsynapsed autosomal regions. Each point is the count from a single cell (total numbers of cells indicated below graphs) from 3 animals of each genotype, except RPA in *Ankrd31^+/−^* (2 mice). Genotype-symbol legend is in panel G. Red lines are mean ± s.d. (I) Time course of γH2AX immunostaining. Arrowheads on pachytene images indicate sex bodies. (J) Reduced γH2AX intensity in *Ankrd31^−/−^* cells. Each point shows the integrated γH2AX immunofluorescent signal from a single spread nucleus. Two independent experiments were conducted, each with a mutant and wild-type pair. For each experiment, values were normalized to the mean from leptonema in wild type. Red lines indicate means (total numbers indicated below the graph). In panels A, D, F, G, H, and J the results of two-tailed Mann Whitney tests are indicated: ns, not significant (p > 0.05), *, p ≤ 0.05; **, p ≤ 0.01; ***, p ≤ 0.001; ****, p ≤ 0.0001. Scale bars throughout are 10 μm except in panel C. See also Figures S3 and S4.

### ANKRD31 deficiency causes dysregulated recombination

Because we isolated ANKRD31 as a REC114 interactor and because the variable synaptic defects in *Ankrd31^−/−^* mutants resemble those in mice with reduced DSB numbers (Kauppi et al., 2013), we hypothesized that ANKRD31 contributes to DSB formation. To test this idea, we examined chromosomal assembly of pre- and post-DSB recombination proteins.

*Ankrd31^−/−^* spermatocytes had fewer and less intense axial foci of REC114 genome-wide throughout prophase I, and REC114 no longer formed blobs on the PAR or other regions associated with the mo-2 minisatellite (**Figures 3D, S3C and S3D**). This functional dependency strengthens the conclusion that ANKRD31 and REC114 interact *in vivo*. MEI1 and MEI4 localization were similarly affected, but not IHO1 (Acquaviva, Jasin & Keeney, unpublished).

To assess recombination, we immunostained chromosome spreads for RAD51 and DMC1 (**Figures 3E–G, S4A and S4B**). In normal meiosis, foci of these proteins appear in leptonema, accumulate to maximal levels in early zygonema, then decline as DSB repair proceeds and autosomes synapse, persisting longer (into mid-pachynema) on the non-homologous portions of the X and Y chromosomes (Moens et al., 2002).

In *Ankrd31^−/−^* spermatocytes, fewer RAD51 and DMC1 foci were present at leptonema (one fourth or fewer of the number in wild type). Foci reached a maximum in early zygonema with a lower than normal average number but with a wide range, such that many cells were similar to wild type (more than 100 foci per cell) while others continued to have substantially fewer foci. As in wild type, focus numbers declined as chromosomes synapsed, but foci accumulated to high levels on unsynapsed axes (**Figure 3E** and “aberrant” class in **Figures 3F and 3G**). Importantly, however, pachytene cells with apparently normal synapsis also displayed elevated numbers of RAD51 and DMC1 foci (**Figures 3F and 3G**), so the increase is not simply because of synaptic errors. Earlier stage cells also had fewer foci of the ssDNA-binding protein RPA (**Figures 3H and S4C**). Seeing fewer RPA foci in early cells suggests that the similar reduction in RAD51 and DMC1 foci is caused by a delay and/or reduced efficiency of forming cytologically observable sites containing resected DSBs, not simply because of defective loading of the strand exchange proteins. Elevated RPA foci were also seen in pachytene cells with normal autosome synapsis, and they accumulated in cells with unsynapsed autosomes (**Figure 3H**). *Ankrd31^+/−^* heterozygotes displayed normal DMC1 and RPA focus numbers (**Figures 3F and 3H**), but had a weak intermediate phenotype for RAD51 foci both at early stages (modest reduction at leptonema and zygonema) and later (modest elevation at pachynema) (**Figure 3G**), possibly indicating a mild haploinsufficient effect.

Reducing DSB numbers to about half the normal level causes pairing and synaptic failure in spermatocytes (Kauppi et al., 2013), so the delayed and reduced numbers of RAD51- and DMC1-associated recombination sites during the critical pairing periods of leptonema and zygonema may explain the partially penetrant synapsis problems in *Ankrd31^−/−^* mutants. The accumulation of foci on unsynapsed axes is a hallmark of homolog engagement failure allowing DSBs to continue forming past the time they normally would have ceased (Kauppi et al., 2013; Thacker et al., 2014).

We also examined phosphorylation of histone variant H2AX (γH2AX). In wild type, γH2AX is formed rapidly by ATM in response to DSBs, then disappears progressively as recombination proceeds until it is largely gone from autosomes in pachynema (**Figure 3I**) (Mahadevaiah et al., 2001; Barchi et al., 2005; Bellani et al., 2005). γH2AX also forms on the X and Y chromosomes as they become transcriptionally silenced and form the sex body (Turner, 2007). *Ankrd31^−/−^* spermatocytes showed pan-nuclear γH2AX staining in leptonema and zygonema, but at lower levels than wild type (**Figures 3I and 3J**). Most of the γH2AX signal dissipated as autosomes synapsed, leaving bright staining in the sex body similar to wild type (**Figure 3I**). Regions of asynapsis retained high levels of γH2AX (**Figure 4A**), as expected (Mahadevaiah et al., 2001). We infer from these patterns that ANKRD31 is dispensable for ATM activation in response to DSBs, but there appears to be less total ATM activity at early stages in keeping with the reduced numbers of DMC1, RAD51, and RPA foci.

**Figure 4.**
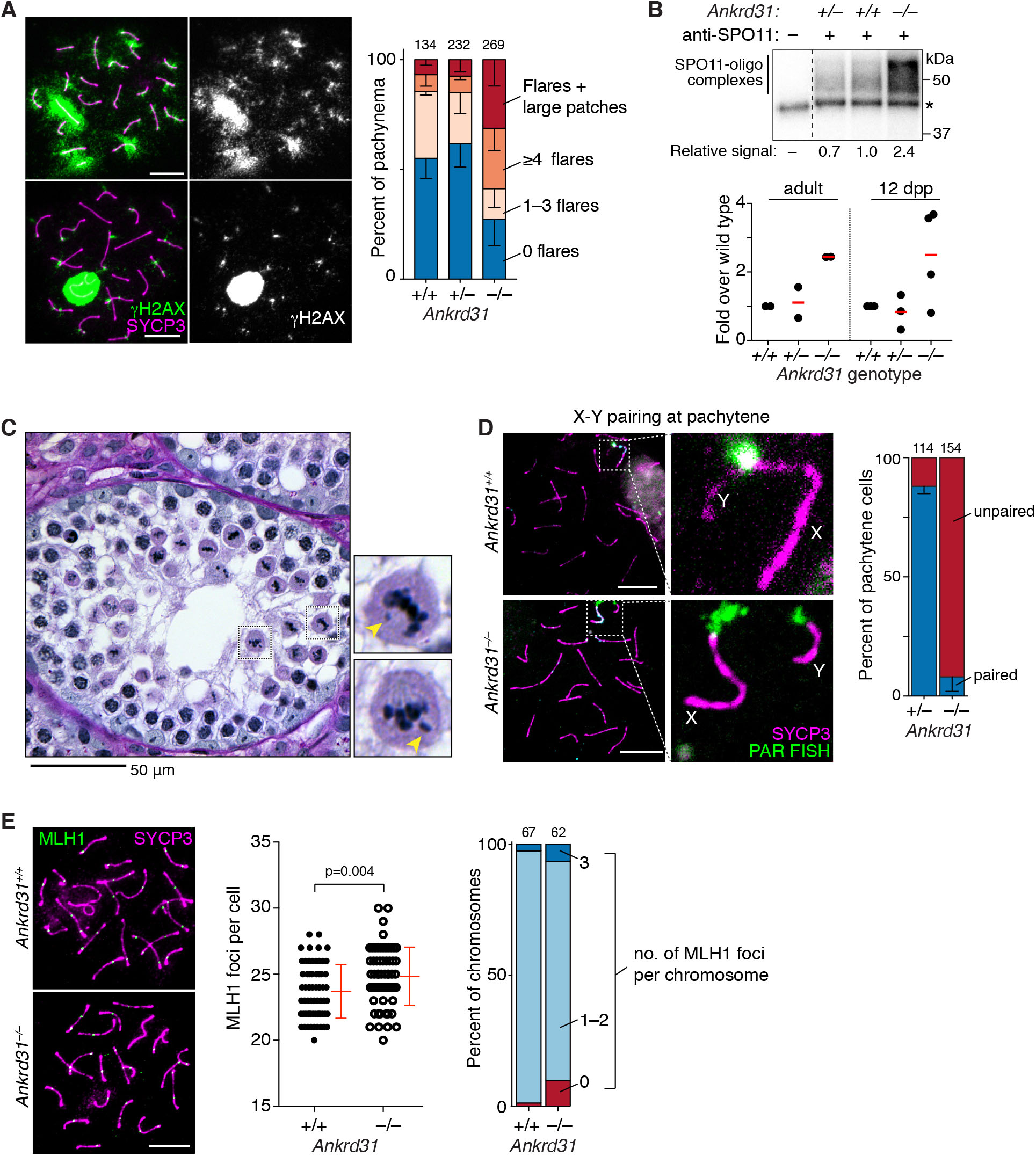
Persistent γH2AX, more DSBs, and metaphase arrest from achiasmate sex chromosomes. (A) Persistent γH2AX. Representative images of *Ankrd31^−/−^* spermatocytes with γH2AX flares and/or larger patches are shown at left. Graph at right quantifies γH2AX patterns from 3 mice of each genotype (total numbers of cells indicated above). Bars indicate mean ± s.d. (B) Increased SPO11-oligo complexes. Upper panel, representative autoradiograph of SPO11-oligo complexes immunoprecipitated from adult testis extracts, radiolabeled with terminal transferase and [α^32^P] dCTP, and separated by SDS-PAGE. Asterisk indicates nonspecific labeling; dashed line indicates where the image was spliced to remove irrelevant lanes. Quantified levels of radioactive signals were background-corrected and normalized to wild type. Graph below summarizes multiple experiments from adults and juveniles. Red bars are means. Levels of SPO11-oligo complexes were significantly different from wild type for *Ankrd31^−/−^* (p = 0.020) but not heterozygotes (p = 0.81) (two-sided one-sample t tests on pooled adult and juvenile data). (C) Representative *Ankrd31^−/−^* seminiferous tubule showing metaphase I spermatocytes with lagging chromosomes (yellow arrowheads). (D) Defective sex chromosome pairing and synapsis. Left, representative pachytene cells immunostained for SYCP3 and subjected to FISH with a PAR-specific probe. Right, quantification of X–Y pairing at pachynema, based on immunofluorescence for SYCP3 and γH2AX, for 3 animals of each genotype (total numbers of cells scored indicated above). Bars indicate mean ± s.d. (E) MLH1 foci. Left, representative pachytene cells. Middle, quantification of MLH1 focus counts per cell from 3 animals of each genotype (p value is from Mann-Whitney test). Right, percent of chromosomes with the indicated number of MLH1 foci (3 animals per genotype; total numbers of cells scored indicated above). Scale bars in B and C are 10 μm. See also Figure S4.

*Ankrd31^−/−^* pachytene spermatocytes with apparently normal autosome synapsis also frequently displayed higher numbers of persistent, discrete flares of γH2AX (**Figure 4A**). This result, along with elevated numbers of DMC1, RAD51, and RPA foci (**Figures 3F–H**), suggests that many cells contain a larger than usual number of incompletely re-paired DSBs. Persistent DSBs at pachynema can trigger spermatocyte apoptosis (Pacheco et al., 2015; Marcet-Ortega et al., 2017), so this may account for some of the partially penetrant pachytene death in *Ankrd31^−/−^* mutants.

Finally, we more directly evaluated DSB formation by labeling SPO11-oligonucleotide complexes, a quantitative by-product of DSB formation and processing (Neale et al., 2005; Lange et al., 2011). Surprisingly, we observed a two to three fold increase in SPO11-oligo complexes in *Ankrd31^−/−^* adults (**Figure 4B**). A similar elevation was observed during the first wave of spermatogenesis in juvenile mice before significant numbers of pachytene cells should be present (**Figures 4B and S4D**), so we can rule out extra DSBs on unsynapsed regions as the source of the increase.

To summarize, recombination foci, γH2AX, and SPO11-oligo complexes demonstrate that DSBs are made, so *Ankrd31^−/−^* does not simply phenocopy *Rec114^−/−^* (Kumar et al., 2018). DSB-associated recombination foci and γH2AX are delayed and reduced, but elevation in SPO11-oligo complexes indicates that more total DNA cleavage events occur. These apparently contradictory findings can be reconciled if ANKRD31 promotes formation of DSB-competent assemblies of REC114 and other proteins, but also assists in local, ATM-dependent downregulation once a DSB has formed (see Discussion). Regardless of the explanation, however, the results clearly demonstrate that DSB formation and recombination are substantially dysregulated in the absence of ANKRD31. Furthermore, the partially penetrant pachytene apoptosis is likely accounted for by a combination of persistent DSBs and synaptic defects.

### Sex chromosome pairing, recombination, and segregation rely on ANKRD31

Metaphase I arrest is typical of mutants that can complete DSB repair but that fail to generate crossovers on some or all chromosomes (Odorisio et al., 1998; Eaker et al., 2002; Kauppi et al., 2011). Indeed, *Ankrd31^−/−^* metaphase I spermatocytes frequently had lagging chromosomes (**Figure 4C**). To determine if specific chromosomes were primarily responsible, we examined spermatocyte spreads. Strikingly, the sex chromosomes were unpaired and asynaptic in 92% of otherwise normal-looking pachytene cells (**Figures 4D and S3A**). By comparison, autosomes showed at most only a modest defect: pachytene cells with complete autosome synapsis had a small increase in the number of MLH1 foci (means of 23.7 in wild type and 24.8 in *Ankrd31^−/−^*; **Figure 4E**), which mark most sites where crossovers will form (Gray and Cohen, 2016). A slight increase in the number of chromosomes lacking an MLH1 focus was seen, but this was not statistically significant (p = 0.105, Fisher’s exact test).

These results indicate that those cells that can progress past pachynema not only successfully navigate pairing and synapsis of autosomes but also efficiently generate autosomal crossovers and chiasmata, thus ensuring proper biorientation on the metaphase I spindle (with perhaps relatively infrequent exceptions). In contrast, the X and Y almost always fail to pair, synapse, and recombine, leading to achiasmate sex chromosomes at metaphase I, which are known to trigger apoptosis (Odorisio et al., 1998; Kauppi et al., 2011). Thus, while ANKRD31 contributes to effective recombination and pairing on autosomes, it is absolutely essential on sex chromosomes in males.

### ANKRD31 shapes the DSB landscape

The data thus far indicated that the sterility of *Ankrd31^−/−^* males was at least partially attributable to downstream consequences of dysregulated DSB formation. Is this strictly an effect on DSB number and timing, or are DSB locations also affected? To answer this question, we mapped DSBs genome wide by sequencing DMC1-bound ssDNA (ssDNA sequencing, or SSDS (Khil et al., 2012)) from testes of adults and 12-dpp juveniles. At the younger age, DSB signal is principally from leptotene and zygotene spermatocytes (Bellve et al., 1977) and thus unaffected by loss of pachytene cells and unlikely to reflect hyper-accumulation of DSBs in unsynapsed regions.

Given the defect in sex chromosome recombination, we examined the PAR and discovered that *Ankrd31^−/−^* mutants display little or none of the high-level SSDS signal normally found there or in PAR-proximal hotspots (**Figure 5A**). This profound defect in forming PAR DSBs (confirmed by cytological quantification of PAR-associated RPA foci (Acquaviva, Jasin & Keeney, unpublished)) is probably sufficient to account for the highly penetrant failures in sex chromosome pairing, synapsis, and recombination.

**Figure 5.**
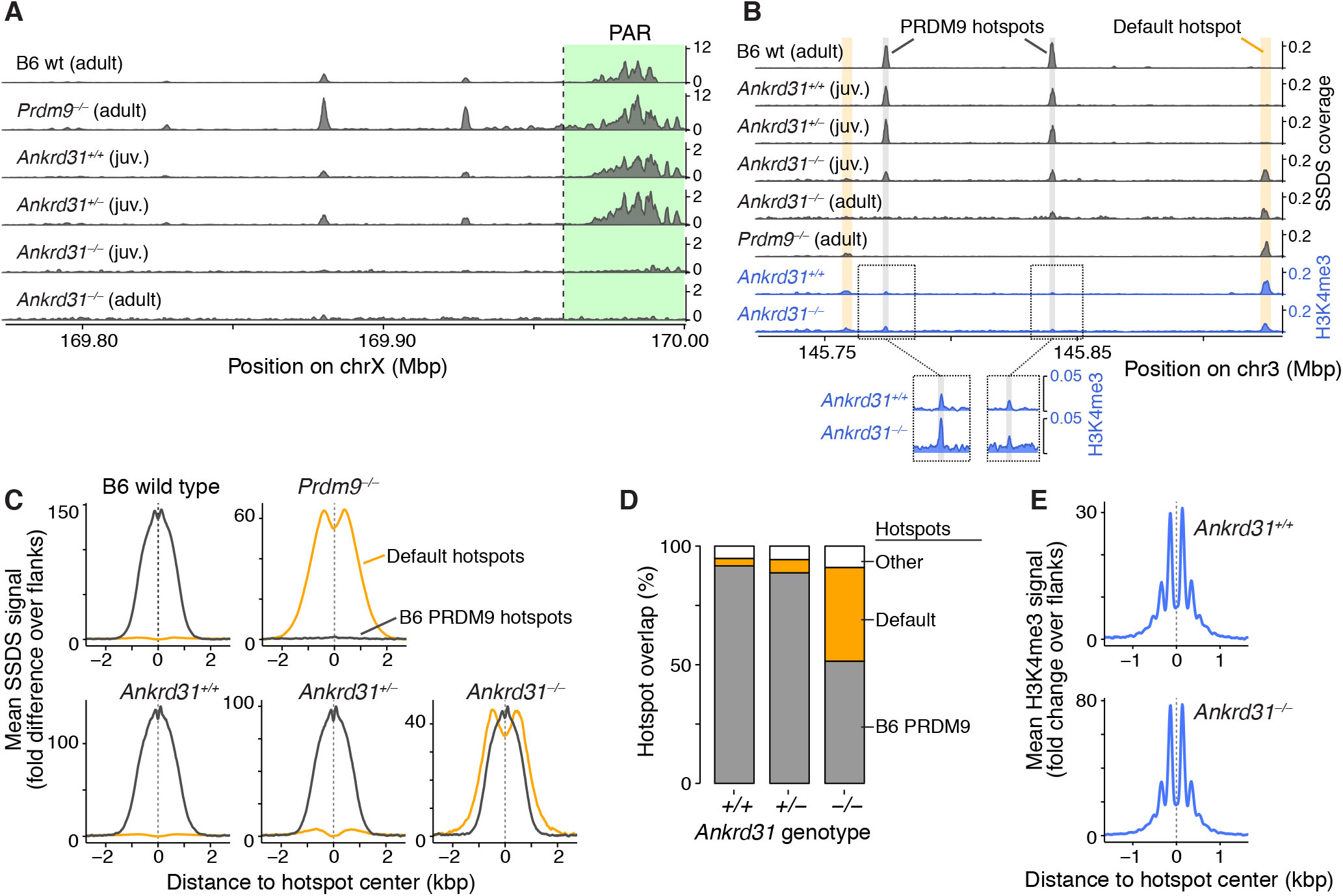
Altered DSB landscape in *Ankrd31^−/−^* males. (A) *Ankrd31^−/−^* mutants are unable to target the PAR and PAR-adjacent hotspots for high-level DSB formation. Browser shots show SSDS maps from adults or juveniles (12 dpp) of the indicated genotypes (smoothed using a 1 kb sliding window in 0.1 kb steps). Only a small portion of the PAR (highlighted green) is present in the mm10 assembly. (B) Browser tracks of an autosomal segment illustrate increased use of default hotspots. Shading highlights a few of the PRDM9-directed (gray) and default (yellow) hotspots; weak hotspots that would be difficult to see at this scale are not highlighted. The bottom two tracks show H3K4me3 ChIP-seq coverage for the same region. Note that H3K4me3 levels tend to be significantly higher at default hotspots than at PRDM9-dependent sites (Brick et al., 2012). (C) Metaplots showing SSDS averages for the indicated genotypes around PRDM9-designated hotspots (black lines, defined in wild-type B6 mice) and default hotspots (yellow lines, defined in *Prdm9^−/−^* mice). (D) Overlap of hotspot calls in SSDS maps from juveniles of the indicated genotype, with either PRDM9-defined (gray) or default (yellow) hotspots. (E) Metaplots showing H3K4me3 ChIP-seq averages around PRDM9-defined hotspot centers. B6 wild-type and *Prdm9^−/−^* SSDS data are from Brick et al. (2012). See also Figure S5.

PAR DSB formation is largely independent of PRDM9 (Brick et al., 2012) (**Figure 5A**) and has other genetic requirements that are distinct from autosomes (Kauppi et al., 2011; Kauppi et al., 2012; Smagulova et al., 2013), so a PAR defect does not necessarily predict that SPO11 targeting to PRDM9 sites would be affected. Surprisingly, however, *Ankrd31*-deficient animals also showed a highly unusual pattern globally: they used many of the PRDM9-targeted hotspots found in B6 mice but also used default (PRDM9-independent) hotspots (**Figures 5B–D**). *Ankrd31^+/−^* heterozygotes appeared largely normal, albeit with slightly increased SSDS signal when averaged over default hotspots, suggesting a subtle haploinsufficient phenotype (**Figure 5C**). Although many default hotspots used in the *Ankrd31^−/−^* mutant were of modest strength, there were also many strong ones (**Figures 5B and S5A**). As a result, 28% of SSDS fragments were in default hotspots, thus more than a quarter of DSBs were redirected away from PRDM9-targeted sites. Furthermore, even the hotspots that were shared often differed in strength in the *Ankrd31^−/−^* mutant (**Figure S5B**).

Mixed use of PRDM9-targeted and default hotspots indicates that ANKRD31 promotes but is not essential for PRDM9 targeting of SPO11 activity. We considered that ANKRD31 might function upstream of PRDM9, i.e., that *Ankrd31^−/−^* mutants have partially lost PRDM9 function. However, *Prdm9* mRNA levels were normal (**Figure S5C**) and, more importantly, H3K4me3 appeared unaffected at both PRDM9-targeted and default sites (**Figures 5B and 5E**). These results imply that PRDM9 expression, binding to chromatin, and histone methyltransferase activity are effectively normal in the absence of ANKRD31. By process of elimination, we suggest that ANKRD31 functions downstream of PRDM9, either directly or indirectly (see Discussion).

Collectively, these findings reveal that ANKRD31 plays a substantial role in shaping the landscape of recombination initiation. In doing so, it has two distinct functions. First, it is essential for preferential DSB formation in PAR, and is thus crucial for sex chromosome recombination in males. Second, it promotes but is not essential for SPO11 cleavage at PRDM9-targeted locations. Thus, paradoxically, absence of ANKRD31 eliminates the use of the special PRDM9-independent sites in the PAR but also increases the use of PRDM9-independent (default) sites globally. Scenarios to reconcile this paradox are described below (see Discussion).

### Structure of an ANKRD31–REC114 complex

To delineate how ANKRD31 and REC114 interact, we characterized them biochemically and structurally. By yeast two-hybrid assay, the N-terminal two-thirds of REC114 (aa 1–147) and the C-terminal tail of ANKRD31 (aa 1810–1857) were necessary and sufficient for interaction (**Figures S6A and S6B**). When expressed in *E. coli*, the N-terminal part of REC114 and the ANKRD31 C-terminal tail (**Figures 6A–C**) also co-purified as a stable complex. We obtained diffraction quality crystals for REC114 aa 1–158 (hereafter REC114_N_) complexed with ANKRD31 aa 1808-1857 (hereafter ANKRD31_C_) and the structure was solved by single-wavelength anomalous diffraction and refined to 2.8 Å resolution (**Table S1**).

**Figure 6.**
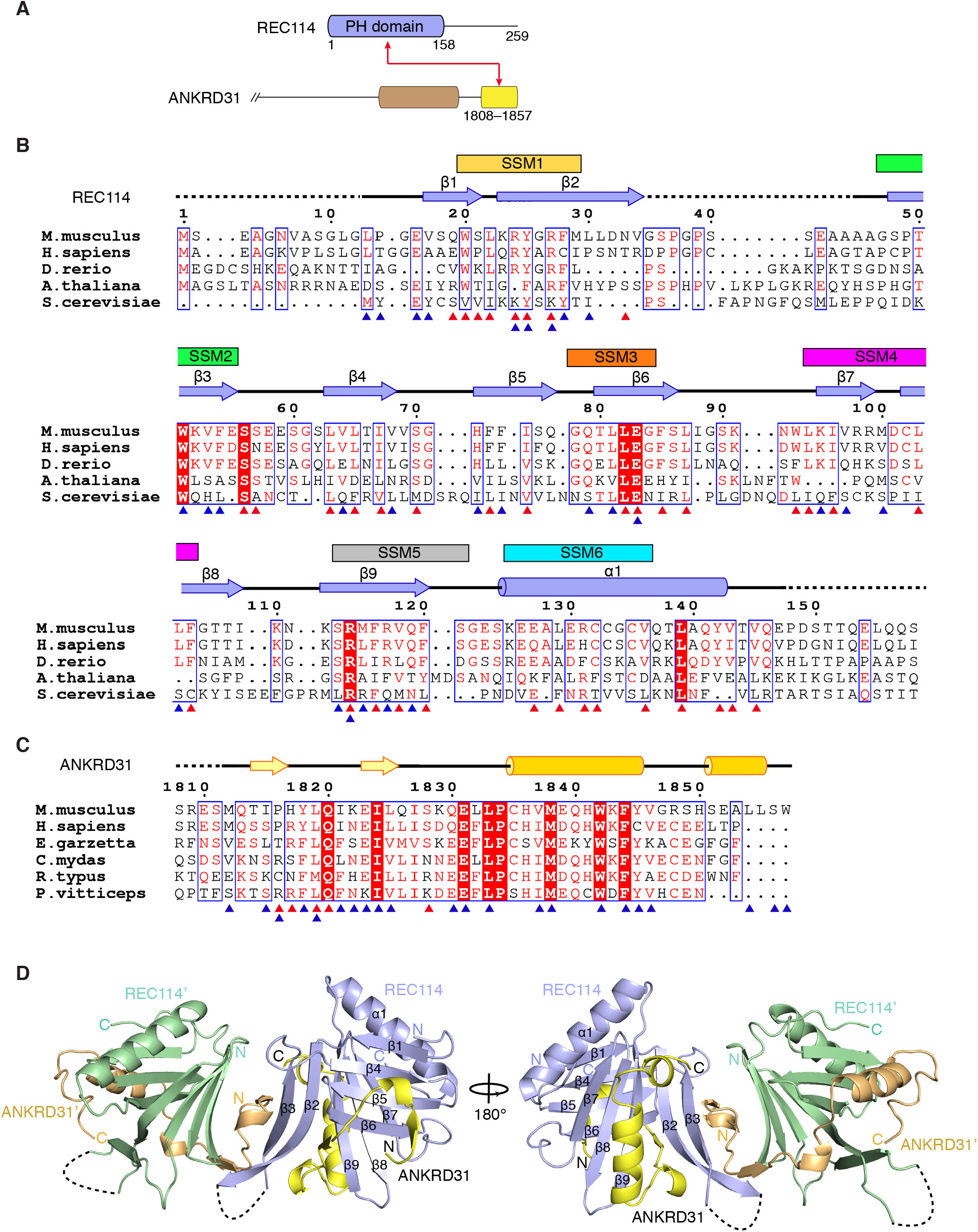
Structure of REC114_N_ in complex with ANKRD31_C_. (A) Domain organization (to scale) of REC114 and C-terminal portion of ANKRD31, with the red arrow identifying segments involved in complex formation. (B and C) Structure-based sequence alignment of REC114_N_ (B) and ANKRD31_C_ (C) orthologs. Secondary structure elements are referenced according to mouse proteins. Invisible residues in the structure are indicated with dashed lines. Red triangles indicate residues involved in intramolecular interactions, blue triangles indicate residues in intermolecular interactions. Boxed residues highlight conservation. The six N-terminal SSMs are mapped above the REC114_N_sequence. Species: *Mus musculus, Homo sapiens, Danio rerio, Arabidopsis thaliana, Saccharomyces cerevisiae, Egretta garzetta, Chelonia mydas, Rhincodon typus, Pogona vitticeps*. (D) Front and back views of the overall structure of a pair of REC114_N_–ANKRD31_C_ heterodimers. See also Figures S6 and S7.

We observed a pair of REC114_N_–ANKRD31_C_ heterodimers in the crystal asymmetric unit (**Figure 6D**). Individual heterodimers superpose well (r.m.s.d. of 0.47 Å **Figure S6C)**, with contacts across heterodimers highlighted in expanded segments of **Figure S6D**. Since these same contacts (**Figure 5D**) are also observed in higher order symmetry units in the crystal lattice (**Figure S6E**), they are assigned to crystal packing interactions. The molecular weights of REC114_N_ alone and of the REC114_N_–ANKRD31_C_ complex, measured by size exclusion chromatography with in-line multi-angle light scattering (SEC-MALS), are comparable with estimates for monomeric REC114_N_ (20.1 kDa measured versus 17.3 kDa predicted) and REC114_N_–ANKRD31_C_ heterodimer (32.9 kDa measured versus 23.3 kDa predicted) **(Figure S5F**), reinforcing that heterodimer contacts are likely due to crystal packing.

REC114_N_ adopts a β-sandwich fold, consisting of two nearly orthogonal antiparallel β-sheets, that is closed at one end by a C-terminal amphipathic α-helix (α1) and remains open at the other end (**Figure 6D**). The N-terminal β-sheet is formed by β1 and β4–β6 and the C-terminal sheet comprises β2, β3, and β7–β9. These secondary structure elements form a hydrophobic core that stabilizes the β-sandwich fold. The loop connecting β2 and β3 is invisible in the density map, indicating high flexibility.

ANKRD31_C_ wraps around the REC114 β-sheets to form an intermolecular sandwich fold, burying ∼1,800 Å^2^ of surface area (**Figure 7A**). Almost all ANKRD31_C_ residues, including the ultimate C terminus, are involved in the interaction with REC114_N_ (**Figures 7A–F**) and can be well traced in the structure. We subdivided ANKRD31_C_ intermolecular contacts within the heterodimer into three segments (S1, S2 and S3), with the impact of interfacial ANKRD31_C_ and REC114_N_ mutations shown in **Figures 7G** and **7H**, respectively.

**Figure 7.**
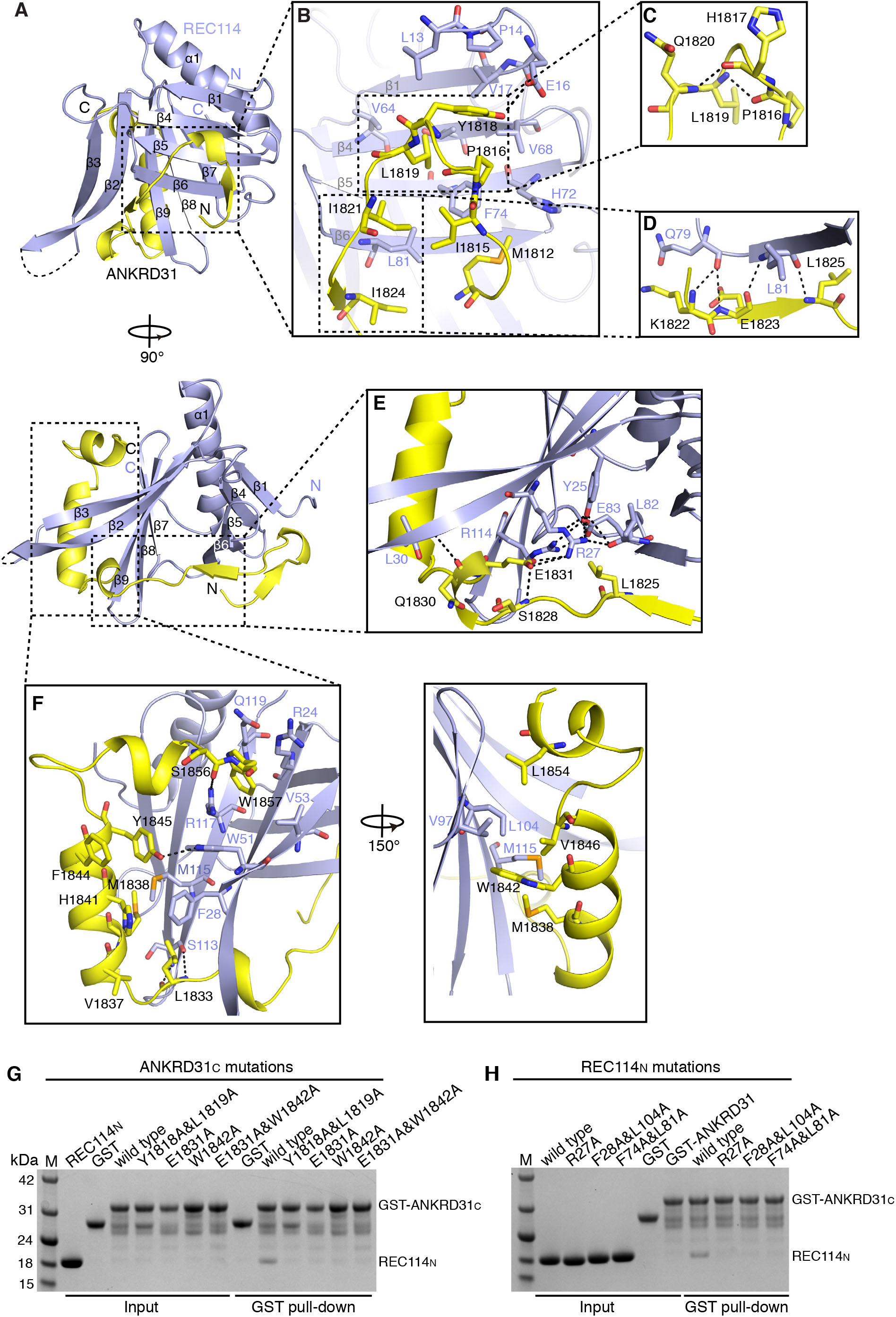
Intermolecular contacts between REC114_N_ and ANKRD31_C_. (A) Front (above) and side (below) views of the REC114_N_–ANKRD31_C_ heterodimer. (B-F) Detailed views of interfaces between ANKRD31_C_ and REC114_N_ in the heterodimeric complex. Hydrogen bonds and salt bridges are shown as black dashed lines. (G, H) Mutational analysis of ANKRD31_C_ (G) and REC114_N_ (H) residues involved in binding, assayed by pull-down assay with GST-tagged ANKRD31_C_. See also Figures S6 and S7.

S1 (aa 1812-1824) forms a turn that strategically positions side chains for mainly hydrophobic contacts with the N-terminal β-sheet of REC114_N_ (**Figure 7B**), with this interaction disrupted by the Y1818A/L1819A double mutation (**Figure 7G**). The importance of the participating hydrophobic residues on the REC114_N_ N-terminal β-sheet was also confirmed by the loss of binding for the dual mutation F74A/L81A (**Figures 7H and S6G**). The turn of S1 is stabilized by two intramolecular main-chain hydrogen bonds formed by P1816-L1819 and H1817-Q1820 segments (**Figure 7C**) and the β-strand at the C terminus of S1 is hydrogen bonded with β6 of REC114_N_ (**Figure 7D**).

S2 (aa 1825-1832) interacts with the open end of the β-sandwich (**Figure 7E**). E1831 is the key residue involved in bridging contacts with REC114_N_, with the E1831 side chain anchored by two salt bridges and one hydrogen bond. The intramolecular hydrogen bond was formed with the main-chain amino group of S1828, and the salt bridges are formed with the side chain of highly conserved R27 of REC114_N_ (**Figures 6B and 7E**). The R27 side chain is further stabilized by a hydrogen-bond and salt bridge network formed by the nearly invariant Y25, L82, E83 and R114 (**Figures 6B and 7E**), which also contributes to the hydrophobic core of the N-terminal and C-terminal β-sheets. Furthermore, the Q1830 main-chain carboxyl group of ANKRD31_C_ hydrogen bonds with L30 from REC114_N_, stabilizing the interaction of E1831 with R27. Mutating ANKRD31_C_ E1831 (E1831A) or REC114_N_ R27 (R27A) dramatically reduced binding (**Figures 7G, 7H and S6G**).

S3 (aa 1833-1857) primarily comprises two amphipathic α-helixes that pack against a relatively hydrophobic groove on the C-terminal β-sheet of REC114_N_ (**Figure 7F**). The importance of these hydrophobic interactions was confirmed by the disruptive effect of the W1842A mutant of ANKRD31_C_ and the F28A/L104A dual mutant of REC114_N_ (**Figures 7G, 7H and S5G**). L1833 forms main-chain hydrogen bonds with S113 of REC114_N_ at the site where ANKRD31_C_ passes through the C-terminal β-sheet between β2 and β9. The side chain of Y1845 hydrogen bonds with strictly conserved W51 of REC114_N_, which stabilizes the conformation of the first α-helix in the S3 segment. The C-terminal W1857 residue inserts into a hydrophobic pocket of REC114_N_formed by the aliphatic side chains of R24, R117, Q119 and V53 (**Figure 7F**), which are strictly conserved in vertebrate REC114 orthologs (**Figure 6B**). This pocket is hydrophilic at the surface but hydrophobic in the interior. The aromatic ring of W1857 also forms a cation-π interaction by packing against the ammonium group of R117 and the aromatic side chain of invariant W51 from REC114_N_. The interaction is further stabilized by the hydrogen bond between S1856 and R117 (**Figure 7F**).

### REC114_N_ has an unexpected pleckstrin homology (PH) domain-like fold

A search of the Protein Data Bank using the Dali server (Holm and Sander, 1995) revealed that REC114_N_ is highly similar to the N-terminal domain of coactivator-associated methyltransferase 1 (CARM1) (Troffer-Charlier et al., 2007), which belongs to the PH domain superfamily. The PH superfamily shares a highly similar fold without significant primary sequence similarity. Structure-based sequence alignment suggested that REC114 orthologs contain similar PH domain-like folds in yeast, plants and animals (**Figure 6B**).

Of seven short signature sequence motifs (SSMs) shared among REC114 orthologs (Maleki et al., 2007; Kumar et al., 2010; Tesse et al., 2017), six can be mapped to our structure and, strikingly, almost all overlap with the secondary structure elements (**Figures 6B and S7A**). We found extra conserved regions from structure-based alignment, although the sequences overall showed substantial divergence. For example, the hydrophobic residues in β4 and β5, together with residues in other secondary structure elements involved in the formation of the hydrophobic core, exhibited significant conservation. By contrast, the variable loops connecting the β-strands have no detectable sequence similarity (**Figure 6B**). Some hydrophilic residues are also highly conserved. S56 at the end of β3 forms hydrogen bonds with L22, K23, S57 and L63, which contributes to the stability at the protein surface (**Figure S7B**). R27, E83 and R114 are involved in both intramolecular hydrogen-bond interactions and intermolecular contacts with ANKRD31_C_ (**Figure 7E**). The R27A mutation eliminated interaction with ANKRD31_C_ (**Figure 7H**).

Interestingly, we also observed conservation of REC114_N_ residues that interact with ANKRD31_C_. For example, invariant W51 contributes to complex formation by hydrogen bonding with Y1845 and cation-π packing with W1857 of ANKRD31_C_ (**Figure 7F**). Other hydrophobic residues on the surface of REC114_N_ involved in the contacts with ANKRD31_C_, including F28, F74, L81, and L104, are also conserved, even in plants and yeast (**Figures 6B and 7H**).

## Discussion

Our search for new components of the mammalian meiotic recombination machinery uncovered mouse ANKRD31 as a direct interaction partner of REC114 and key regulator of multiple aspects of recombination initiation. The principles underlying molecular recognition were elucidated from a crystal structure of the REC114_N_–ANKRD31_C_ complex, with observed intermolecular contacts validated from mutational analysis of interfacial residues from both components. ANKRD31 is required for normal assembly of REC114 complexes on chromosomes and for normal location, timing and number of DSBs genome wide and especially in the PAR. Presumably as a consequence of defects in these critical functions, absence of ANKRD31 causes catastrophic failure of spermatogenesis and greatly diminished oogenesis. Studies of this possibly vertebrate-specific factor expand the catalog of essential mammalian meiotic proteins and elucidate how cells control SPO11-generated DSBs. Our results agree well with independent findings of Papanikos et al. (A. Tóth, personal communication).

### Paradoxical mutant phenotypes shed light on ANKRD31 functions

*Ankrd31^−/−^* mutants presented two apparent paradoxes: 1) There were fewer/delayed sites marked by recombination protein foci but also more total DSBs as measured by SPO11-oligo complexes. 2) There was increased use of some PRDM9-independent sites (default sites genome-wide) but also failure to use others in and near the PAR. These seemingly contradictory findings can be parsimoniously reconciled and provide important insights into the mechanism and regulation of DSB formation.

A key to the first paradox is recognizing that cytological foci report on events at low resolution (>100-nm scale), whereas SPO11-oligo complexes report at finer biochemical scale. A simple way to reconcile our findings would be if individual DMC1, RAD51, and RPA foci in *Ankrd31^−/−^* mutants often contain clusters of two or more DSBs in close proximity. Such DSB clusters occur occasionally in wild-type yeast and more frequently in mutants lacking the ATM ortholog Tel1 (Garcia et al., 2015). Because of the reduced number and intensity of REC114 foci, we favor the interpretation that *Ankrd31^−/−^* mutants make fewer and less efficient DSB-forming machines and thus have delayed and/or fewer recombination sites, but we further propose that these machines are also less tightly controlled and often generate multiple DSBs. In this model, ANKRD31 is a lynchpin of DSB regulation with both DSB-promoting and -suppressing activities. A nonexclusive alternative explanation for the reduced recombination foci would be defects in DSB resection, which generates the ssDNA to which RPA, DMC1, and RAD51 bind (Mimitou et al., 2017). We cannot rule out this possibility, but it would not explain the increase in SPO11-oligo complexes, diminished REC114 assemblies, or reduced γH2AX levels.

*Ankrd31^−/−^* mutants partially phenocopy *Atm^−/−^* for increased SPO11-oligo complexes (Lange et al., 2011; Pacheco et al., 2015), so we speculate that ANKRD31 contributes to ATM-dependent inhibition of DSBs. ANKRD31 may do so as a direct ATM target because it has 24 SQ/TQ sites, the phosphorylation motif for ATM and related kinases (Traven and Heierhorst, 2005). It is also possible that ANKRD31 acts upstream of ATM by promoting kinase activation by DSBs. The reduced γH2AX formation could reflect such a role in ATM activation, or could simply reflect delayed and reduced numbers of sites where ATM is being activated.

For the second paradox, we propose that ANKRD31 has distinct roles in targeting SPO11 activity to different parts of the genome, but that these roles are related through recruitment of REC114. We found that ANKRD31 colocalizes with REC114 in large assemblies in the PAR and in smaller foci along unsynapsed axes genome wide. The large REC114 assemblies are eliminated in the absence of ANKRD31, whereas the smaller foci are still present but become fainter and less numerous. In separate work, we found that the PAR undergoes dramatic structural reorganization during meiosis—the chromosome axes elongate and sister axes split apart—and that these behaviors require ANKRD31-dependent accumulation of REC114, MEI4, and other pro-DSB factors (Acquaviva, Jasin & Keeney, unpublished). That study also showed that ANKRD31 and MEI4 are mutually dependent for their accumulation on the PAR (Acquaviva, Jasin & Keeney, unpublished). The absolute requirement for ANKRD31 in the PAR could mean that ANKRD31 is a primary determinant of this localization, i.e., that it recognizes a cis-acting feature of the PAR itself and recruits REC114 (and possibly other proteins) by direct interaction. In this model, we hypothesize that the ability to recruit REC114 has allowed ANKRD31 to acquire a highly specialized function in ensuring high-level DSB formation in the PAR. PRDM9 happens to be largely or completely irrelevant for this targeting, at least in mice (Brick et al., 2012; Hinch et al., 2014).

Elsewhere in the genome, where PRDM9 is the major arbiter of hotspot positions, ANKRD31 is important but not essential for normal targeting of SPO11 activity. Since our data suggest that PRDM9 itself is able to function more or less normally in *Ankrd31^−/−^* mutants, one possibility is that ANKRD31 directly links SPO11 to PRDM9-established marks, a process that remains poorly understood (Imai et al., 2017; Parvanov et al., 2017; Diagouraga et al., 2018; Grey et al., 2018). An alternative possibility is suggested by the interpretation above that the delay in accumulation of DMC1 foci is because there are fewer and less efficient DSB-forming machines. Perhaps the PRDM9-dependent histone marks have either vanished or become unusable for other reasons by the time the later DSBs are being made. In this model, the direct function of ANKRD31 is to stabilize association of REC114 and other proteins on chromosomes to promote efficient DSB formation, with effects on PRDM9 targeting an indirect consequence. Regardless of the underlying mechanism, however, our results definitively establish that the two modes of PRDM9-independent targeting (default hotspots vs. the PAR) are mechanistically distinct because they respond in opposite ways to ANKRD31 deficiency.

### ANKRD31 in male and female fertility

*Ankrd31^−/−^* males displayed an unusual spermatogenic arrest phenotype, partially penetrant in pachytene and more fully penetrant in metaphase I. The latter arrest is likely attributable to rampant failure of sex chromosome recombination tied to PAR dysfunction, but achiasmate autosomes may also contribute. The pachytene arrest appears to trace its origins to a mixture of problems. Specifically, we propose that delayed and reduced DSB formation during a critical window early in prophase (leptonema through early zygonema) leads to stochastic defects in homologous pairing and synapsis (Kauppi et al., 2013). Altered DSB locations and formation of clustered DSBs may also contribute to asynapsis. Cells with synaptic defects die in pachytene from a recombination checkpoint, sex body failure, or both (Royo et al., 2010; Marcet-Ortega et al., 2017). We further surmise that some pachytene cells die from persistent DSBs (evidenced by γH2AX flares and recombination protein foci) even though they achieve full synapsis. This apparent repair defect could be an indirect consequence of dysregulated DSB formation, e.g., from cutting of both sister chromatids, and/or could mean that ANKRD31 has a repair-promoting function after the DSB is made.

Oocytes do not make sex bodies and therefore do not have the same arrest behavior as spermatocytes in the face of synaptic defects (Hunter, 2017). Moreover, because the two X chromosomes can recombine at any point along their lengths, they do not rely on the PAR for pairing and chiasma formation. However, oocytes do eliminate cells with persistent DSBs, typically as cells transition to dictyate arrest and form follicles (Di Giacomo et al., 2005; Bolcun-Filas et al., 2014; Hunter, 2017). A plausible hypothesis, therefore, is that similar molecular defects contribute to oocyte loss in *Ankrd31^−/−^* females as cause the partially penetrant pachytene cell death in males. If so, then the sexually dimorphic *Ankrd31^−/−^* phenotype tracks with known differences between oocytes and spermatocytes for both quality control surveillance and for the uniquely challenging constraints on sex chromosome recombination in males.

### ANKRD31 as a modular scaffold to facilitate and regulate DSB formation

The unexpected similarity of REC114_N_ to PH domains (independently demonstrated by a crystal structure of REC114_N_ alone (Kumar et al., 2018)) provides a clear explanation for previously recognized patterns of sequence conservation in REC114 orthologs (Maleki et al., 2007; Kumar et al., 2010; Tesse et al., 2017) by demonstrating that most of the conserved SSMs correspond to critical secondary structure elements. Many especially well conserved residues contribute to the folding of the REC114 PH domain, but others are surface exposed. Some of the latter mediate interactions with ANKRD31_C_ but other conserved residues that are not part of this intermolecular interface are likely to mediate interactions with other proteins and/or with the REC114 C-terminal region.

Yeast two-hybrid assays and characterization of recombinant proteins demonstrated that the ANKRD31–REC114 interaction is direct, and crystallographic analysis delineated at atomic resolution how they interact. The ANKRD31_C_ peptide wraps around REC114_N_ and makes extensive contacts, resulting in a stable heterodimer. The ANKRD31_C_sequence and REC114-interacting residues therein are highly conserved in vertebrate ANKRD31 orthologs, suggesting evolutionary conservation of the interaction with REC114. Perhaps more intriguing, however, many of the residues in REC114_N_ that interact with ANKRD31_C_ are also highly conserved, even in species such as *A. thaliana* and *S. cerevisiae* that do not have an obvious ANKRD31 ortholog. We therefore speculate either that ANKRD31 is more widely conserved than currently apparent, or that ANKRD31 takes advantage of a REC114 surface that is evolutionarily constrained because it binds a different partner(s) in other species (and perhaps also in competition with ANKRD31 in vertebrates).

We are intrigued by the presence of additional conserved domains in ANKRD31, including the two ARDs and the two conserved domains (CD3 and CD4) in between ARD-2 and ANKRD31_C_. ARDs are often involved in protein-protein interactions, including direct interactions with nucleosomes (Saredi et al., 2016), so we consider it likely that the large ANKRD31 protein acts as a modular scaffold that anchors REC114 to other proteins and to meiotic chromosomes.

## Materials and Methods

Further information and requests for reagents may be directed to Scott Keeney at Memorial Sloan Kettering Cancer Center. All primer sequences are listed in **Supplemental Table S2**. Antibodies are listed in **Supplemental Table S3**.

### Mice

Mice were maintained and sacrificed under U.S.A. federal regulatory standards and experiments were approved by the Memorial Sloan Kettering Cancer Center (MSKCC) Institutional Animal Care and Use Committee. Animals were fed regular rodent chow with ad libitum access to food and water.

Endonuclease mediated alleles (*em1, em2*) were generated by the MSKCC Mouse Genetics core facility by targeting exon 3. Guide RNA with sequence (5ʹ - GTTGTTGCTGGCTCTTAGTG) was cloned into pU6T7 vector (Romanienko et al., 2016). In vitro-transcribed guide RNA (100 ng/μl) and Cas9 (50 ng/μl) were microinjected using conventional techniques (Romanienko et al., 2016) into pronuclei of CBA/J × C57BL/6J F2 hybrid zygotes generated by crossing CBAB6F1/J hybrid females with C57BL/6J males. Genomic DNA from founder animals was amplified with primers ANKRD31-A and ANKRD31-B and digested with T7 endonuclease I to identify potential indel-carrying animals.

To determine the spectrum of mutant alleles in T7-positive *Ankrd31^em^* founder mice, the targeted region was amplified by PCR of tail-tip DNA (ANKRD31-A and ANKRD31-B primers) and sequenced on the Illumina MiSeq platform (Illumina Inc, San Diego, California) at the MSKCC Integrated Genomics Operation. Reads were aligned to mouse genome assembly GRCm37/mm9 and variants were identified using Genome Analysis Toolkit version 2.8-1-g932cd3a (McKenna et al., 2010; DePristo et al., 2011; Van der Auwera et al., 2013). Variants with a minimum variant frequency of 0.01 were annotated using VarScan v2.3.7 software (Koboldt et al., 2012).

Founder mice were crossed to C57BL/J6 mice purchased from Jackson laboratories to obtain germline transmission, then heterozygous animals were backcrossed to C57BL/J6 a further ≥3 generations. Experimental animals were generated by crossing *Ankrd31^+/−^* heterozygous males with either *Ankrd31^+/−^* heterozygous or *Ankr31^−/−^* homozygous females. Genotyping was performed by PCR on genomic DNA with primers 16G008 and 16G011, followed with digestion by AflII (NEB) which recognizes *em1* (+A) allele or with HrpChIV (NEB), which recognizes *em*2 (+C) allele. The PCR product is 339 bp and restriction digestion products are 264 and 75 bp. The systematic names of the alleles generated in this study are *Ankrd31^em1Sky^* and *Ankrd31^em2Sky^*.

### Yeast two-hybrid screen

Mouse *Mei4* and *Rec114* cDNA were amplified by PCR and cloned into pGBKT7 vector (Clontech). Briefly, PCR products were amplified from testis cDNA library and digested by EcoRI (NEB) and NdeI (NEB) and then cloned into linearized pGBKT7. pGBKT7-*Mei4* or pGBKT7-*Rec114* plasmids were transformed into Y2HGold yeast two-hybrid strain following manufacturer’s instructions (Clontech, 630439). Single colonies were picked to verify protein expression by western blotting and to ensure that neither pGBKT7-*Mei4* nor pGBKT7-*Rec114* auto-activated Gal4-driven reporters promoters or were toxic to yeast.

The yeast two-hybrid library was prepared from cDNA generated from whole-testis extracts of 12-dpp males using the Make Your Own “Mate & Plate” Library System according to the manufacturer’s instructions (Clontech, 630490). We estimated that the library contained ~10^6^ independent clones.

The yeast-two hybrid screen was performed using Matchmaker Gold Yeast Two-Hybrid System according to the manufacturer’s instructions (Clontech, 630489). Briefly, individual clones of Y2H bait strains containing pGBKT7-*Mei4* or pGBKT7-*Rec114* were inoculated in liquid synthetic dextrose medium lacking tryptophan (SD-Trp) and grown to OD_600_ of ~0.8, then cells were pelleted and resuspended in SD-Trp at 10^8^ cells/ml. Bait strains were mixed with aliquots of the library strain and incubated in 2× YPDA medium with 50 μg/ml kanamycin for 20–24 hours at 30°C, shaking at 40 rpm. Cultures were checked for the presence of zygotes by light microscopy, and then cells were plated on 50 plates (150 mm × 15 mm) of SD lacking tryptophan and leucine and containing X-α-galactose and aureobasidin A (SD-Trp/Leu/X-α-gal/AbA), and incubated for 5 days at 30°C. Approximately 10^6^ zygotes were screened for each bait. Those that formed a blue colony were re-streaked on fresh SD-Trp/Leu/X-α-gal/AbA plates. If also positive in this second growth test, they were subjected to PCR amplification as instructed in Matchmaker Insert Check PCR Mix 2 (Clontech, 630497).

The plasmids were then recovered from yeast and transformed into *E. coli* and sequenced, using Easy Yeast Plasmid Isolation Kit (Clontech, 630467). The screen with pGBKT7-*Mei4* as bait yielded clones containing sequences from *Rec114* and *Med10*. The screen with pGBKT7-*Rec114* yielded *Ankrd31*, *Sohlh1*, *Coro1b*, *Phka2*, *Rps20*, *Sin3b*, *Ctnna1*, and *Hsp90ab*.

### Targeted yeast two-hybrid assays

Genes were amplified from cDNA generated from testes of 12-dpp C57Bl/6J mice as described above. Genes of interest were amplified and cloned into either pGADT7 or pGBKT7. Mating of bait- and prey-containing strains (Y187 and Y2HGold yeast strains) and selection on SD-Trp/Leu/X-α-gal/AbA plates were performed following manufacturer’s instructions as described above (Clontech). Primers used for cloning are listed in **Supplemental Table S2**.

### Analysis of sequence conservation

Amino acid sequence divergence (fraction of residues changed) and Ka and Ks values were downloaded from HomoloGene Release 68 (April 2014), https://www.ncbi.nlm.nih.gov/homologene. Estimated times to last common ancestor were obtained from http://www.timetree.org/ (Hedges et al., 2015) (accessed July 26, 2015).

For each HomoloGene entry with mammalian orthologs (n = 19,498) we generated a least-squares linear regression line fitting amino acid sequence divergence as a function of time since last common ancestor, forcing an intercept of zero. The slope of this line (multiplied by 100) is the divergence rate (percent amino acid changes per Myr) plotted in **Figure S1B**. We also calculated the Ka/Ks ratio for each pairwise comparison between mammalian species within a HomoloGene entry, and took the median value among these as the representative value for that HomoloGene entry. This is the median Ka/Ks ratio plotted in **Figure S1B**.

Likely ANKRD31 orthologs were identified by BLAST searches using full-length human ANKRD31 as the query, or using just the C-terminal 70 amino acids. From a collection of over 150 homologs including representatives from mammals, reptiles, and birds, we generated a multiple sequence alignment of the C-terminal 70 amino acids using MUSCLE (Edgar, 2004) in MegAlign Pro (Lasergene v 14.1.0), then used this alignment as input for a HHMER3 search (http://hmmer.org/). This search again identified ANKRD31 homologs in vertebrates, including one in *Gasterosteus aculeatus* (three-spined stickleback; UniProt G3N8J6), but failed to find any significant hits in fungi. When the stickleback protein was used as a BLAST query, we identified additional homologs from fish that had not turned up in the earlier searches. Interestingly, the fish homologs all have only one ARD, which by multiple sequence alignment appeared to be a better match for ARD-2 in the mammalian proteins.

### Antibody generation

To raise polyclonal antibodies against ANKRD31, a fragment of *Ankrd31* coding sequence (corresponding to a.a. 1–324) was cloned into pETDuet™-1 (Millipore) using the In-Fusion cloning kit (Clontech, Takara). This expression vector was used to produce 6×His-ANKRD31_1–134_. Briefly, *E. coli* strain BL21 was transformed with the vector and a single colony was picked to inoculate LB overnight at 37°C. The overnight culture was diluted to ~0.04 (OD_600_) and once OD_600_ had reached ~0.6, the protein expression was induced by addition of 1 mM Isopropyl β-D-1-thiogalactopyranoside (IPTG) for 2 hrs at 37°C. Cells were pelleted by centrifugation at 6000 g, 15 min. Pellets were resuspended in lysis buffer (50 mM HEPES-NaOH pH 8.0, 300 mM NaCl, 0.1 mM dithiothreitol (DTT), 10% glycerol and 20 mM imidazole) and stored at −80°C. To purify the ANKRD31 fragment, the pellet was thawed at room temperature and placed on ice and all further steps were executed at 4°C. The sample was sonicated 8 times 15 seconds on-off, 20% duty cycle. The solution was cleared by centrifugation at 19,000 rpm, 1 hr. Equilibrated Ni-NTA Agarose (QIAGEN, 30230) was added to the cleared protein lysate and incubated for 1 hr, then the lysate-resin slurry was transferred into Poly-prep chromatography columns (BioRad, 731-1550) and washed with lysis buffer several times. The protein was eluted with elution buffer (lysis buffer + 500 mM imidazole) and fractions containing the ANKRD31 fragment were combined and subjected to size-exclusion chromatography. Sample was loaded on a Superdex 200 column equilibrated with 50 mM HEPES-NaOH pH 8.0, 300 mM NaCl, 1 mM DTT, 10% glycerol and 5 mM EDTA and the fractions under the main peak were collected and analyzed on SDS-PAGE. Purified protein fractions were combined and concentrated (Millipore), then frozen in liquid nitrogen and stored at −80°C. Purified protein fragment was used to immunize two rabbits and two guinea pigs by Pocono Rabbit Farm & Laboratory. Polyclonal anti-ANKRD31 antibodies were enriched from serum using NAb™ Protein A Plus Spin Columns (Thermoscientific, 89952). Polyclonal antibody specificity was tested by immunofluorescent staining of meiotic chromosome spreads and immunoprecipitation followed by western blotting.

### Total mRNA extraction, cDNA library generation and RT-qPCR

Testes tissue from *Ankrd31^+/+^* and *Ankrd31^−/−^* animals were dissected and frozen on dry ice. Total mRNA was extracted using RNeasy Plus Mini Kit (QIAGEN, 74134) following the manufacturer’s instructions. Superscript™ III First-Strand Synthesis SuperMix (Invitrogen, 18080400) was used with oligo dT primers to generate testis cDNA, which was diluted 1:10 to be used in RT-qPCR carried out using LightCycler 480 SYBR Green I Master (Roche, 4707516001). Amplification products were detected on the LightCycler 480 II Real-Time PCR instrument (Roche). LightCycler 480 Software was used to quantify products by absolute quantification analysis using the second derivative maximum method. All reactions were done in triplicate and the mean of crossing point (Cp) value were used for the analysis. Cp values were normalized to the value obtained for *B2M* reactions (ΔCp). Then, the differences between knockout and wild-type samples were calculated for each primer set (ΔΔCp) and the fold change (knockout vs wild type) was calculated as 2^−ΔΔCp^. Primers used for RT-qPCR are listed in Supplemental Table S2.

### Immunoprecipitation and western blot analysis of ANKRD31

Dissected testes were placed in an Eppendorf tube and snap frozen on liquid nitrogen and stored at -−80°C. The frozen tissue was resuspended in RIPA buffer (50 mM Tris-HCl pH 7.5, 150 mM NaCl, 0.1% SDS, 0.5% sodium deoxycholate, 1% NP40) supplemented with protease inhibitors (Roche Mini tablets). The tissue was disrupted with a plastic pestle. The homogenized extract was supplemented with 10 mM MgCl_2_ and Benzonase nuclease (EMD Millipore (70664-3), 28 unit/μl) and incubated with end-over-end rotation for 1 hr at 4°C. The samples were centrifuged at 15,000 rpm for 20 min. The clear lysate was transferred to a new tube and used for the immunoprecipitation. Whole-cell extract was pre-cleared with protein A/G Dynabeads (Thermofisher, 1004D, 1001D) by end-over-end rotation for 1 hr at 4°C. Antibodies (home-made, rabbit anti-ANKRD31 or guinea pig anti-ANKRD31, 1–2 μg) were added to pre-cleared lysates and incubated overnight with end-over-end rotation at 4°C. Protein A/G Dynabeads were added to the tubes and incubated for 1 hr with end-over-end rotation, at 4°C. Beads were washed three times with RIPA buffer, resuspended in 1× NuPAGE LDS sample buffer (Invitrogen) with 50 mM DTT, and incubated 10 min at 70°C to elute immunoprecipitated proteins.

For western blotting, samples were separated on 3–8% Tris-Acetate Nu-PAGE precast gels (Life Technologies) at 150 V for 70 min. Proteins were transferred to polyvinylidene difluoride (PVDF) membranes by wet transfer method in Tris-Glycine-20% methanol, at 120 V for 40 min at 4°C. Membranes were blocked with 5% non-fat milk in 1× phosphate buffered saline (PBS)-0.1% Tween (PBS-T) for 30 min at room temperature on an orbital shaker. Blocked membranes were incubated with primary antibodies (guinea pig anti-ANKRD31, 1:4000) 1 hr at room temperature or overnight at 4°C. Membranes were washed with PBS-T for 30 min at room temperature, then incubated with HRP-conjugated secondary antibodies (rabbit anti-guinea pig IgG (Abcam), 1:10,000) for 1 hr at room temperature. Membranes were washed with PBS-T for 15 min and the signal was developed by ECL Prime (GE Healthcare).

### Spermatocyte chromosome spreads

Testes were dissected and deposited after removal of the tunica albuginea in 50 ml Falcon tubes containing 2 ml TIM buffer (104 mM NaCl, 45 mM KCl, 1.2 mM MgSO_4_, 0.6 mM KH_2_PO_4_, 6.0 mM sodium lactate, 1.0 mM sodium pyruvate, 0.1% glucose). 200 μl collagenase (20 mg/ml in TIM buffer) was added, and left shaking at 550 rpm for 55 min at 32°C. Every 15 min, tissue was checked and carefully pulled further apart to enhance digestion. After incubation, TIM buffer was added to a final volume of 15 ml, followed by centrifugation for 1 min at 600 rpm at room temperature. Supernatant was decanted and this wash and centrifuge procedure was repeated 3 times. Separated tubules were resuspended in 2 ml TIM, with 200 μl trypsin (7 mg/ml in TIM) and 20 μl DNase I (400 μg/ml in TIM buffer) and incubated for 15 min. at 32°C at 550 rpm in a thermomixer. 500 μl trypsin inhibitor (20 mg/ml in TIM) and 50 μl DNase I solution were added and mixed. A wide mouthed plastic Pasteur pipette was used to disperse the tissue further by pipetting up and down for 2 min. The Pasteur pipette was first used to pipet a 2% BSA solution in PBS to coat the plastic and minimize cell loss. Cells were passed through a 70-μm cell strainer into a new 50 ml Falcon tube. TIM was added to a final volume of 15 ml and mixed. Cells were centrifuged for 5 min at 1200 rpm. Supernatant was decanted, 15 µl DNase I solution was added and gently mixed, followed by 15 ml TIM. Washing with TIM and resuspension in presence of DNase I was repeated 3 times. Single-cell suspension was pelleted and resuspended in TIM according to original weight (~200 mg in 400 μl). 10 μl of cell suspension was added to 90 μl of 75 mM sucrose solution, flicked three times and incubated for 8 min at room temperature. Superfrost glass slides were divided in two squares by use of Immedge pen, each square received 100 μl 1% paraformaldehyde (PFA) (freshly dissolved in presence of NaOH at 65°C, 0.15% Triton, pH 9.3, cleared through 0.22 μm filter) and 45 μl of cell suspension was added per square, swirled three times, and dried in a closed slide box for 3 hr, followed by drying with half-open lid 1.5 hr at room temperature. Slides were washed in a Coplin jar 2 × 3 min in milli-Q water on a shaker, 1 × 5 min with 0.4% PhotoFlow, air dried and stored in aluminum foil in –80°C.

### Immunostaining

Slides of spread spermatocytes were blocked for 30 min at room temperature in 100 ml solution containing 1× PBS with 0.05% Tween-20 and 3 mg/ml bovine serum albumin (BSA). Slides were incubated with primary antibody overnight in a humid chamber at 4°C. Slides were washed 3 × 10 min in PBS– 0.05% Tween, then incubated with secondary antibody 45 min at 37°C in a humid chamber. Slides were washed 3 × 5 min in the dark on a shaker with PBS– 0.05% Tween, before air drying and mounting with Vectashield containing DAPI. All primary and secondary antibodies used here are listed in **Supplemental Table S3**.

### PAR FISH

All following steps were performed in the dark to prevent loss of fluorescent signal. After regular staining for immunofluorescence, slides were re-fixed in 2% PFA in PBS for 10 min at room temperature. Slides were rinsed once in PBS, and washed for 4 min in PBS at room temperature. Slides were washed with 70% ethanol for 4 min, followed by 4 min wash with 90% ethanol, and final wash with 100% ethanol for 5 min. Slides were air dried vertically for 5 min. Slides were incubated with 15 μl of PAR probe BAC RP24-500I4 and coverslips were sealed with rubber cement (Weldwood contact cement). Slides were denatured for 7 min at 80°C, followed by overnight incubation (>14 hr) in a humid chamber at 37°C. Coverslips were removed, and slides were rinsed in 0.1× SSC buffer, washed in 0.4× SSC, 0.3% NP-40 for 5 min, followed by a wash in PBS–0.05% Tween-20. Slides were washed once in H_2_O, dried and mounted in Hardset Vectashield (without DAPI).

### Expansion microscopy

Expansion microscopy was performed by the MSKCC Molecular Cytology Core Facility based on published methods (Tillberg et al., 2016). Slides of spread spermatocyte chromosomes were prepared and stained as described above. Slides were treated with Acryloyl X-SE (AcX) (0.05 mg/ml diluted in PBS) overnight at room temperature. Samples were washed twice for 15 min with PBS, followed by treatment with monomer solution (8.6% (w/v) sodium acrylate, 2.5% (w/v) acrylamide, 0.1% (w/v) N, Nʹ-methylenebisacrylamide, 11.7% (w/v) NaCl in PBS) for 30 min at 4°C. Gelation solution was prepared from chilled monomer solution on ice by mixing in the following order 4-hydroxy-2,2,6,6-tetramethylpiperidin-1-oxyl (4-hydroxy-TEMPO) inhibitor solution (final concentration 0.01% (w/v)), 10% tetramethylethylenediamine TEMED (final concentration 0.2% (w/v)). 10% APS solution (final concentration 0.2% (w/v)) was added to initiate the gelling process. The solution was briefly vortexed, and the sample was placed in imaging spacer, used as a well on a glass slide coated with SigmaCoat. The well was sealed with a cover glass and incubated for 2 hr at 37°C in a humidified chamber. Gelled samples were transferred to a 6-well glass bottom plate and treated with digestion buffer (50 mM Tris-HCl pH 8.0, 23 mM EDTA, 0.5% Triton X-100, 0.8 M guanidine HCl) containing freshly added proteinase K (final concentration 200 μg/ml). The sample was digested for 3 hr at 60°C, or until the gel detached from the slide. The container was sealed to prevent drying out during digestion. Digestion buffer was removed and replaced with PBS, sample was stained with DAPI for 30 min at room temperature. DAPI solution was removed and sample was washed in excess of dH_2_O, three times, 30 min each to expand the sample. Water was removed, and the sample was embedded in 2% agarose in dH_2_O. Samples were imaged after agarose was fully solidified.

### Histology

Fixation, tissue processing, and staining were performed as described (Jain et al., 2018). Testes from adult mice were fixed overnight in 4% PFA at 4°C, or in Bouin’s fixative for 4 to 5 hr at room temperature. Bouin’s fixed testes were washed in water for 1 hr at room temperature, followed by five 1-hr washes in 70% ethanol at 4°C. Wild-type and mutant ovaries were fixed in 4% PFA overnight at 4°C. PFA-fixed tissues were washed 4 × 5 minutes in water at room temperature. Fixed tissues were stored in 70% ethanol for up to 5 days prior to embedding in paraffin and sectioning (5 μm for testes, 8 μm for ovaries). The tissue sections were deparaffinized with EZPrep buffer (Ventana Medical Systems) and antigen retrieval was performed with CC1 buffer (Ventana Medical Systems). Sections were blocked for 30 min with Background Buster solution (Innovex), followed by avidin-biotin blocking for 8 min (Ventana Medical Systems). Periodic acid Schiff (PAS) staining and immunohistochemical TUNEL assay were performed by the MSKCC Molecular Cytology Core Facility using the Autostainer XL (Leica Microsystems, Wetzlar, Germany) automated stainer for PAS with hematoxylin counterstain, and using the Discovery XT processor (Ventana Medical Systems, Oro Valley, Arizona) for TUNEL. Ovaries were fixed with 4% PFA overnight and serially sectioned at 8 μm. Sections were incubated with anti-VASA for 5 hr, followed by 60 min incubation with biotinylated goat anti- rabbit (Vector Labs, cat# PK6101) at 1:200 dilution. The detection was performed with DAB detection kit (Ventana Medical Systems) according to manufacturer’s instruction. Slides were counterstained with hematoxylin and coverslips were mounted with Permount (Fisher Scientific).

### Histological examination of somatic tissues

Gross histopathological analysis of major organs and tissues was performed by the MSKCC Laboratory of Comparative Pathology for the following male mice: two *Ankrd31* mutants and two heterozygous littermates aged 28 weeks. The following females were analyzed: one *Ankrd31* mutant and one wild-type littermate aged 24 weeks; one *Ankrd31* mutant and one heterozygous littermate aged 15 weeks. Histological examination of the following tissues was performed: diaphragm, skeletal muscle, sciatic nerve, heart/aorta, thymus, lung, kidneys, salivary gland, mesenteric lymph nodes, stomach, duodenum, pancreas, jejunum, ileum, cecum, colon, adrenals, liver, gallbladder, spleen, uterus, ovaries, cervix, urinary bladder, skin of dorsum and subjacent brown fat, skin of ventrum and adjacent mammary gland, thyroid, parathyroid, esophagus, trachea, stifle, sternum, coronal sections of head/brain, vertebrae and spinal cord. Tissues were fixed in 10% neutral buffered formalin and bones were decalcified in formic acid solution using the Surgipath Decalcifier I (Leica Biosystems, Wetzlar, Germany) for 48 hr. Samples were routinely processed in alcohol and xylene, embedded in paraffin, sectioned (5 μm), and stained with hematoxylin and eosin. Mutant males examined had marked degeneration of seminiferous tubules with aspermia. In heterozygous littermates testes appeared normal, but some minimal germ cell exfoliation in seminiferous tubules was observed, and minimal intraluminal cell debris in epididymides, which were considered likely to be naturally occurring for this age and genetic background. All follicle types were absent in the ovaries of the 24 week old homozygous knockout; the mutant female of 15 weeks appeared to have a reduced number of oocytes and follicles (not quantified). Some obesity associated conditions were observed, such as moderate hepatic lipidosis, but these were not attributable to *Ankrd31* genotype. All other findings in mutants were considered incidental and/or age-related.

### Image acquisition

Images of spread spermatocytes were acquired on a Zeiss Axio Observer Z1 Marianas Workstation, equipped with an ORCA-Flash 4.0 camera, illuminated by an X-Cite 120 PC-Q lightsource, with either 63× 1.4 NA oil immersion objective or 100× 1.4 NA oil immersion objective. Marianas Slidebook (Intelligent Imaging Innovations, Denver Colorado) software was used for acquisition.

Imaging of the expanded sample was performed on a Zeiss LSM 880 confocal microscope with a 63× 1.4 NA oil immersion objective. Z-stacks were acquired with 488 and 561 laser lines used for excitation. Zeiss Airyscan detector has been used for imaging to increase the resolution of the 4× expanded sample further. All imaging data was acquired at optimal pixel sizes and optimal axial intervals. Data were processed using the Zen software (Carl Zeiss, Jena, Germany).

Whole slides (histology), either PAS or TUNEL stained, were scanned and digitized with the Panoramic Flash Slide Scanner (3DHistech, Budapest, Hungary) with a 20× 0.8 NA objective (Carl Zeiss, Jena, Germany). High resolution images of PAS and IHC images were acquired with a Zeiss Axio Imager microscope using a 63× 1.4 NA oil immersion objective (Carl Zeiss, Jena, Germany).

### Quantification of SPO11-oligo complexes

We purified SPO11 oligos from testes of adults (>8 weeks of age) or 12-dpp juvenile mice as previously described (Lange et al., 2011; Pan et al., 2011). Testis were decapsulated, flash frozen in liquid nitrogen and stored at –80°C. Testes were homogenized with prechilled plastic pestles in lysis buffer (1% Triton X-100, 25 mM HEPES-NaOH pH 7.4, 5 mM EDTA) containing EDTA-free protease inhibitors, cleared by centrifugation for 25 min at 4°C, 100,000 rpm, followed by transfer of supernatant to low binding Eppendorf tubes. Supernatants were incubated with mouse anti-SPO11-180 antibody (3 μg per immuno-precipitation (IP) sample) before protein A-agarose beads (1 hr 4°C end-over-end rotation) were added for an additional 3 hr at 4°C. SPO11 IP samples were eluted in Laemmli sample buffer for 4 min at 95°C. Beads from the SPO11 IP were washed 3 times with IP buffer (1% Triton X-100, 150 mM NaCl, 15 mM Tris-HCl pH 8.0), and washing steps were consolidated into eluates. Combined IP eluate and washes were diluted with 5 volumes of IP buffer before performing a second round of IP (SPO11 IP2). Anti-Mouse SPO11-180 antibody was added (3 μg per sample) for 1 hr at 4°C with end-over-end rotation, before addition of protein A-agarose beads and further mixing overnight. SPO11 IP2 samples were washed 3 times with IP wash buffer and 2 times with 1× NEB4 before radio-labeling with terminal deoxynucleotidyl transferase and [α-^32^P] CTP in NEB buffer 4 for 1 hr at 37°C in a thermomixer at 300 rpm. Beads were washed 5 times with IP buffer before elution with 2× Laemmli sample buffer. Proteins were separated by SDS-PAGE on 8% Invitrogen BOLT gels, 125 V for 90 min. Proteins were transferred to PVDF membrane via semi-dry transfer in 210 mM Tris, 103 mM glycine, 0.04% SDS, pH 9.5) for 17 min at 17 V. Radiolabelled species were detected and quantified after 48 hr exposure with Fuji phosphor screens and quantified in ImageJ.

### SSDS and H3K4me3 ChIP

DMC1-SSDS was performed as described (Brick et al., 2018) except that lysis was performed in RIPA buffer (10 mM Tris-HCl pH 8, 1 mM EDTA, 0.5 mM EGTA, 1% Triton X-100, 0.1% sodium deoxycholate, 0.1 % SDS, 140 mM NaCl) instead of lysis buffer containing 1% SDS. Therefore, the dialysis step was omitted. 24 μg of DMC1 antibody (C-20, sc-8973, discontinued) was used to pull down the nucleoprotein filaments.

H3K4me3 ChIP was performed as described (Brick et al., 2012) except that MNase treatment was substituted for sonication. Testes were dissected and placed in cold PBS, then were moved to 10 cm tissue culture dish (Gibco), tunica albuginea was removed and testes were placed on a rocker in a 15 ml Falcon tube containing 10 ml 1% freshly made paraformaldehyde (PFA) in MilliQ H_2_O for 10 min at room temperature. One ml of 1.25 M glycine was added for 5 min after which samples were put on ice. Fixed tissue was homogenized in a 15 ml Dounce homogenizer by 10 strokes with the tight pestle. Samples were filtered through a 70-μm cell strainer into a 15 ml Falcon tube, centrifuged for 5 min at 4°C at 900 g. The supernatant was removed, the pellet resuspended in 10 ml PBS and centrifuged at 900 g for 5 min at 4°C. Pellets were resuspended in 10 ml hypotonic lysis buffer (HLB) (10 mM Tris-HCl pH 8.0, 1 mM KCl, 1.5 mM MgCl_2_ and 250 μl 40 Protease inhibitor (Roche 11836170001)). Samples were left on rocker for 30 min at 4°C, followed by homogenization with 15 ml Dounce homogenizer. Samples were centrifuged for 10 min at 10,000 g at 4°C. The pellet containing nuclei was resuspended in 1 ml MNase buffer (50 mM Tris-HCl pH 8.0, 1 mM CaCl_2_, 4 mM MgCl_2_, 4% NP-40) and 27 μl 40x protease inhibitor. Samples were transferred to low protein binding Eppendorf tubes (#022431081). MNase (150 units per sample) was added (USB affymetrix, J70196-ZCR) and incubated for 5 min at 37°C. 21 μl 0.5 M EDTA was added (final concentration 10 mM) and incubated for 5 min at 4°C. Samples were spun down at 15,000 rpm in table top centrifuge at 4°C for 10 min. Supernatant was transferred to new Eppendorf tubes and centrifuged again. Supernatant was split into two identical volumes in new tubes and diluted 1:3 with RIPA buffer (10 mM Tris-HCl pH 8, 1 mM EDTA, 0.5 mM EGTA, 1% Triton X-100, 0.1% sodium deoxycholate, 0.1 % SDS, 140 mM NaCl). 10 μg of H3K4me3 antibody (Abcam, ab8580) was added per sample (5 μg per tube) and incubated with end-over-end rotation over-night at 4°C. 100 μl of Protein G Dynabeads per tube were aliquoted and washed 3× with RIPA buffer using magnetic holder. Beads were pelleted, resuspended in 50 μl of digested chromatin sample and transferred to tube with the rest of the chromatin sample, followed by 2 hrs incubation at 4°C with end-over-end rotation. Beads were collected using a magnetic holder, the supernatant removed, and beads were subject to washes with 1.2 ml of each of the following buffers in order: low salt immune complex wash buffer (0.1% SDS, 1% Triton X-100, 2 mM EDTA, 20 mM Tris-HCl pH 8.0, 150 mM NaCl), high salt immune complex wash buffer (0.1% SDS, 1% Triton X-100, 2 mM EDTA, 20 mM Tris-HCl pH 8.0, 500 mM NaCl), LiCl immune complex wash buffer (0.25 M LiCl, 1% IGEPAL-CA630, 0.5% sodium deoxycholate, 1mM EDTA, 10 mM Tris-HCl pH 8.0), and 2 washes of TE buffer (10 mM Tris-HCl, 1 mM EDTA pH 7.4). Beads were pelleted and supernatant removed. 150 μl of elution buffer (1% SDS, 100 mM NaHCO_3_ in MilliQ water) was added and beads were flicked, before incubation at 65°C for 30 min, shaking in a thermomixer at 500 rpm. Beads were pelleted, supernatant was transferred to new tube, in which previously split samples were now combined. 12 μl of 5 M NaCl was added to each sample and incubated overnight (18–20 hr) at 65°C. Each sample received 6 μl of 0.5 M EDTA, 12 μl of 1 M Tris-HCl pH 6.5 and 5 μl of proteinase K (20 mg/ml) followed by 2 hr incubation at 45°C. Samples were cleaned up using the Qiagen MinElute column following manufacturer instructions, with the alteration that samples were diluted with 7 volumes of PB buffer and passed over columns at a volume of 700 μl, each time with 1 min binding and 1 min centrifugation until all sample had been applied. Columns were washed with 750 μl PE buffer, and samples were finally eluted using 12 μl EB buffer. Immunoprecipitated DNA was diluted to 50 μl using Ultrapure H_2_O and was quantified by PicoGreen and the size was evaluated on a High Sensitivity BioAnalyzer chip. When possible, fragments between 100 and 600 bp were size selected using a MPure XP beads (Beckman Coulter catalog # A63882) and Illumina libraries were prepared using the KAPA Hyper Prep Kit (Kapa Biosystems KK8504) according to the manufacturer’s instructions with 8-10 cycles of PCR. Barcoded libraries were sequenced on a Hiseq 2500 in rapid mode in a 50 bp x 50 bp paired-end run, using the HiSeq Rapid SBS Kit v2 (Illumina). An average of 59 million paired reads was generated per sample.

### Protein expression and purification for crystallography

The mouse ANKRD31 (residues 1808-1857, ANKRD31_C_) and REC114 (residues 1-158, REC114_N_) were cloned into a modified RSFduet-1 vector (Novagen) with an N-terminal His_6_-SUMO tag on ANKRD31_C_ and no tag on REC114_N_. ANKRD31_C_ and REC114_N_ were co-expressed in *E. coli* strain BL21(DE3) RIL (Stratagene). The cells were grown at 37°C until OD_600_ reached 0.8, then the media was cooled to 16°C and IPTG was added to a final concentration of 0.35 mM to induce protein expression overnight at 16°C. The cells were harvested by centrifuge at 4°C and disrupted by sonication in Binding buffer (20 mM Tris-HCl pH 8.0, 500 mM NaCl, 20 mM imidazole) supplemented with 1 mM PMSF (phenylmethylsulfonyl fluoride) and 3 mM β-mercaptoethanol. After centrifugation, the supernatant containing complexes of ANKRD31_C_ and REC114_N_ was loaded onto 5 ml HisTrap Fastflow column (GE Healthcare). After extensive washing with Binding buffer, the complex was eluted with Binding buffer supplemented with 500 mM imidazole. The His_6_-SUMO tag was removed by Ulp1 protease digestion during dialysis against Binding buffer and separated by reloading onto HisTrap column. The flow-through fraction was further purified by HiTrap Q FF column and Superdex 75 16/60 column (GE Healthcare). The pooled fractions were concentrated to 5 mg/ml in crystallization buffer (20 mM Tris-HCl pH 8.0, 300 mM NaCl, 1 mM DTT). For the selenomethionine (SeMet) derivative protein, the cells were grown in M9 medium supplemented with lysine, threonine, phenylalanine, leucine, isoleucine, valine, and Se-methionine. GST-tagged ANKRD31_C_ proteins also have C-terminal His_6_ tag and were first purified by HisTrap Fastflow column. Mutants were generated by the QuikChange Mutagenesis Kit (Stratagene). The SeMet derivative protein, GST-tagged protein and all the mutants were purified as described above.

### Crystallization, data collection and structure determination

Crystals of SeMet derivative REC114_N_-ANKRD31_C_ complex were grown from a solution containing 0.1 M Bis-Tris pH 5.5, 25% PEG 3350 using the hanging drop vapor diffusion method at 20°C. For data collection, the crystals were flash frozen (100 K) and collected on NE-CAT beam line 24ID-C at the Advanced Photo Source (APS), Argonne National Laboratory. The diffraction data were processed with HKL2000 (Otwinowski et al., 2003). The structure of REC114_N_-ANKRD31_C_ complex was solved by the single-wavelength anomalous diffraction method using PHENIX (Adams et al., 2002). The automatic model building was carried out using the program PHENIX AutoBuild (Adams et al., 2002). The resulting model was completed manually using COOT (Emsley et al., 2010) and PHENIX refinement (Adams et al., 2002). The statistics of the diffraction and refinement data are summarized in **Table S1**. All molecular graphics were generated with the PyMOL program (https://pymol.org/2/). The sequence alignments were generated with Multalin (Corpet, 1988) and ESPript (Robert and Gouet, 2014).

### GST pull-down assay

50 ng GST-tagged ANKRD31_C_ and mutants were incubated with 50 μL Glutathione Sepharose 4B beads (GE heathcare) at 4°C for 4 h. The beads were washed two times with 1 ml crystallization buffer and then 100 ng of pre-purified REC114_N_ and mutants were added and incubated for 2 h at 4°C. An additional three washes were applied and each sample was analyzed with SDS-PAGE.

### SEC-MALS

SEC-MALS experiments were performed by using an Äkta-MALS system. Proteins (500 μl) at a concentration of 2.5 mg/ml were loaded on Superdex 75 10/300 GL column (GE Healthcare) and eluted with crystallization buffer at a flow rate of 0.3 ml/min. The light scattering was monitored by a miniDAWN TREOS system (Wyatt Technologies) and concentration was measured by an Optilab T-rEX differential refractometer (Wyatt Technologies). Molecular masses of proteins were analyzed using the Astra program (Wyatt Technologies).

### Statistical analysis

Statistical analyses were performed using R version 3.4.4 and RStudio version 1.1.442 (R Core Team, 2018), and Graphpad Prism 7. Statistical parameters and tests are reported in the figures and corresponding figure legends. Sample size, statistical test used, means and error bars are defined on figures and/or in figure legends. No statistical tests was used to predetermine sample size. One of the experiments quantifying SPO11-oligo complexes in 12 dpp animals was performed blinded as to genotype of the animals. No data from experiments were excluded from analysis. In cases where outliers were removed for plotting purposes, none of the data points were removed from the statistical analysis.

### Image analysis

Single cells were manually cropped and analyzed with semi-automated scripts in Fiji (Schindelin et al., 2012). Scripts are available on Github: https://github.com/Boekhout/ImageJScripts. Briefly, for detection of number of foci, images were auto-thresholded on SYCP3 staining, which was used as a mask to use ‘Find Maxima’ to determine number of RPA, DMC1 or RAD51 foci. Images were manually inspected to determine that there were no obvious defects in determining SYCP3 axis, axis from neighboring cells were counted, artifacts were present or clear foci were missed by the script. For detection of ANKRD31 and REC114 foci, overlap and intensity with SYCP3 was detected with a thresholded algorithm regardless of meiotic stage, and a difference of Gaussian (DoG) blurs was used to isolate REC114 or ANKRD31 foci. All images were manually inspected upon analysis.

For γH2AX intensity analysis, leptotene and early zygotene cells were identified by SYCP3 staining, regions of interest were manually drawn and used to measure both the integrated density of a cell of interest, and for a region containing no cells for background subtraction. Integrated density was normalized to mean integrated intensity of wild type leptotene cells per experiment to combine different experiments.

### Data and software availability

The atomic coordinates have been deposited in the Protein Data Bank (PDB) (accession pending). SSDS and H3K4me3 ChIP-seq sequencing data have been deposited in the Gene Expression Omnibus (GEO) repository under accession number GSE118913. Image analysis scripts are available on Github: https://github.com/Boekhout/ImageJScripts

## End Matter

### Author Contributions and Notes

MB characterized male and female phenotypes of *Anrd31^−/−^* mice; performed H3K4me3 ChIP and provided samples for SSDS; identified minimal interacting domains in REC114 and ANKRD31 and generated the first interaction-defective mutants; and wrote the scripts for semiautomated analysis of axis-associated DSB protein foci. MEK performed the yeast two-hybrid screen and identified, cloned, and characterized expression of *Ankrd31*; generated *Ankrd31* mutant mice and performed preliminary characterization; generated and validated anti-ANKRD31 antibodies; and generated and purified the recombinant REC114_N_-ANKRD31_C_ complex. JW refined expression and purification of the REC114_N_-ANKRD31_C_ complex, grew crystals, and performed the X-ray crystallographic analysis, SEC-MALS assays, and GST pull-down experiments on REC114_N_ and ANKRD31_C_ mutants under the supervision of DJP. LA performed FISH experiments and contributed unpublished data and analysis concerning the PAR defects and behaviors of REC114 and other DSB proteins. DYE generated constructs and performed yeast two-hybrid analysis under supervision of MB. FP and KB performed SSDS and analyzed the SSDS and H3K4me3 data under the supervision of RDC. SK conceived the project, analyzed data, performed the homology analysis, and oversaw the research. DJP and JW wrote the structure sections with input from SK. SK wrote the rest of the paper with input from MB, MEK, FP, KB, and RDC. All authors edited the manuscript.

The authors declare no conflict of interest.

## Acknowledgments

We thank Attila Tóth and Bernard de Massy for discussions and sharing unpublished information. We thank Keeney lab members C. Claeys Bouuaert, N. Mohibullah, M. van Overbeek, and S. Kim for experimental advice. We thank MSKCC core facilities, supported by NIH cancer center core grant P30 CA008748: Mouse Genetics (P. Romanienko and W. Mark) for generating the *Ankrd31* knockout; Molecular Cytology (K. Manova) for histology and expansion microscopy; the Integrated Genome Operation (A. Viale and N. Mohibullah) for sequencing; Bioinformatics (N. Socci) for analysis; and the Laboratory of Comparative Pathology (S. Monette) for necropsy. This work utilized the computational resources of the NIH HPC Biowulf cluster (http://hpc.nih.gov). X-ray diffraction studies were conducted at the Advanced Photon Source on the Northeastern Collaborative Access Team beamlines, which are supported by NIGMS grant P41 GM103403 and U.S. Department of Energy grant DE-AC02-06CH11357. The Pilatus 6M detector on 24-ID-C beamline is funded by a NIHORIP HEI grant (S10 RR029205). MB was supported in part by a Rubicon fellowship from the Netherlands Organization for Scientific Research. LA was supported in part by a fellowship from the Lalor Foundation. This work was supported in part by NIGMS grant R35 GM118092 (SK), the Howard Hughes Medical Institute (SK), and funds from the Maloris Foundation (DJP).

---

**Figure S1.**
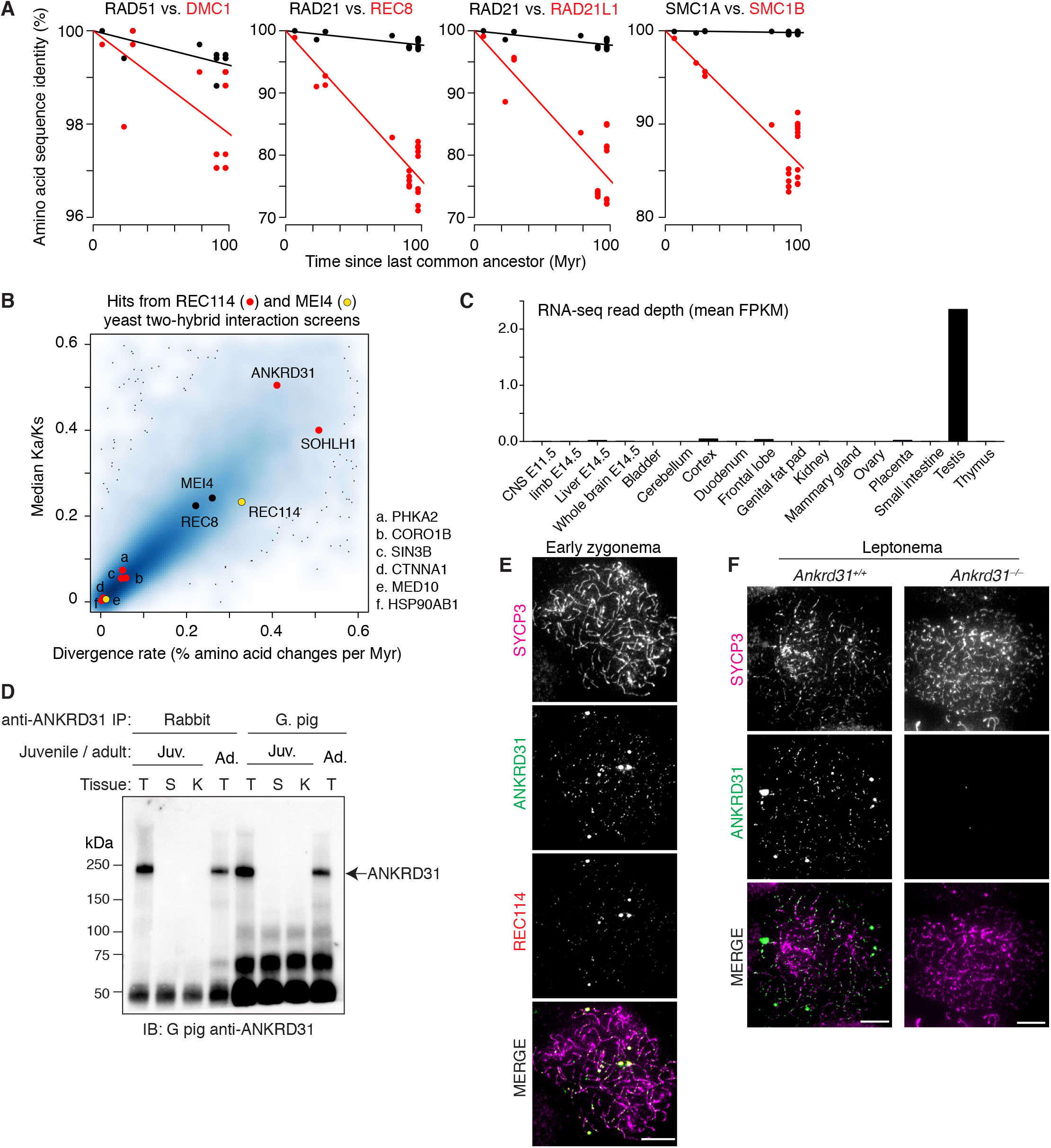
Identification and characterization of *Ankrd31*, related to Figure 1. (A) Meiosis-specific proteins typically show more rapid sequence divergence than their more ubiquitously expressed paralogs. Graphs show amino acid sequence identity as a function of time since last common ancestor for pairwise comparisons of orthologs in different mammalian species for either a meiosis-specific protein (red) or its ubiquitously expressed paralog (black). Species included: *Homo sapiens, Pan troglodytes, Maccaca mulatta, Canis familiaris, Bos taurus, Mus musculus, and Rattus norvegicus*. Lines are least squares fits with intercepts set at 100% identity. Note differences in y axes: the trend of more rapid divergence of the meiotic paralog is true regardless of overall rate of sequence divergence for the specific protein type. Even for the relatively highly conserved proteins DMC1 and RAD51, the meiotic version is more rapidly diverging. (B) Plots of Ka/Ks (ratio of the number of nonsynonymous substitutions per non-synonymous site to the number of synonymous substitutions per synonymous site) vs. amino acid sequence divergence rate within mammals (see Methods). Data for all proteins in the NCBI HomoloGene database (Release 68 (April 2014), https://www.ncbi.nlm.nih.gov/homologene) were plotted using the smoothScatter function in R (blue to gray shading), and overlaid with hits from the yeast two-hybrid screens (red and yellow points). MEI4 and REC8 are shown for comparison. Among the REC114 two-hybrid hits, ANKRD31 and SOHLH1 stood out as having relatively high divergence rates and high Ka/Ks ratios. SOHLH1 is a basic helix-loop-helix transcription factor that regulates germ cell development (Ballow et al., 2006). It is expressed in germline stem cells before meiosis and controls spermatogonial and oocyte differentiation without apparent direct involvement in meiosis per se (Ballow et al., 2006; Shin et al., 2017). We therefore did not explore potential links of SOHLH1 to REC114 function, and instead focused on ANKRD31. (C) *Ankrd31* RNA-seq read depth in various tissues (FPKM, fragments per kb per million reads). (D) Immunoprecipitation (IP) and immunoblotting (IB) of ANKRD31 from extracts of testis (T), spleen (S) or kidney (K) from 17-dpp juveniles or testis from adult, using antisera raised in the indicated species. (E) ANKRD31 localizes to chromatin. A wild-type spermatocyte at early zygonema is shown (intermediate stage between spermatocytes shown in Figure 1F). (F) Specificity of the guinea pig anti-ANKRD31 antiserum. Representative spreads are shown at equivalent exposures. Scale bars in E, F are 10 μm.

**Figure S2.**
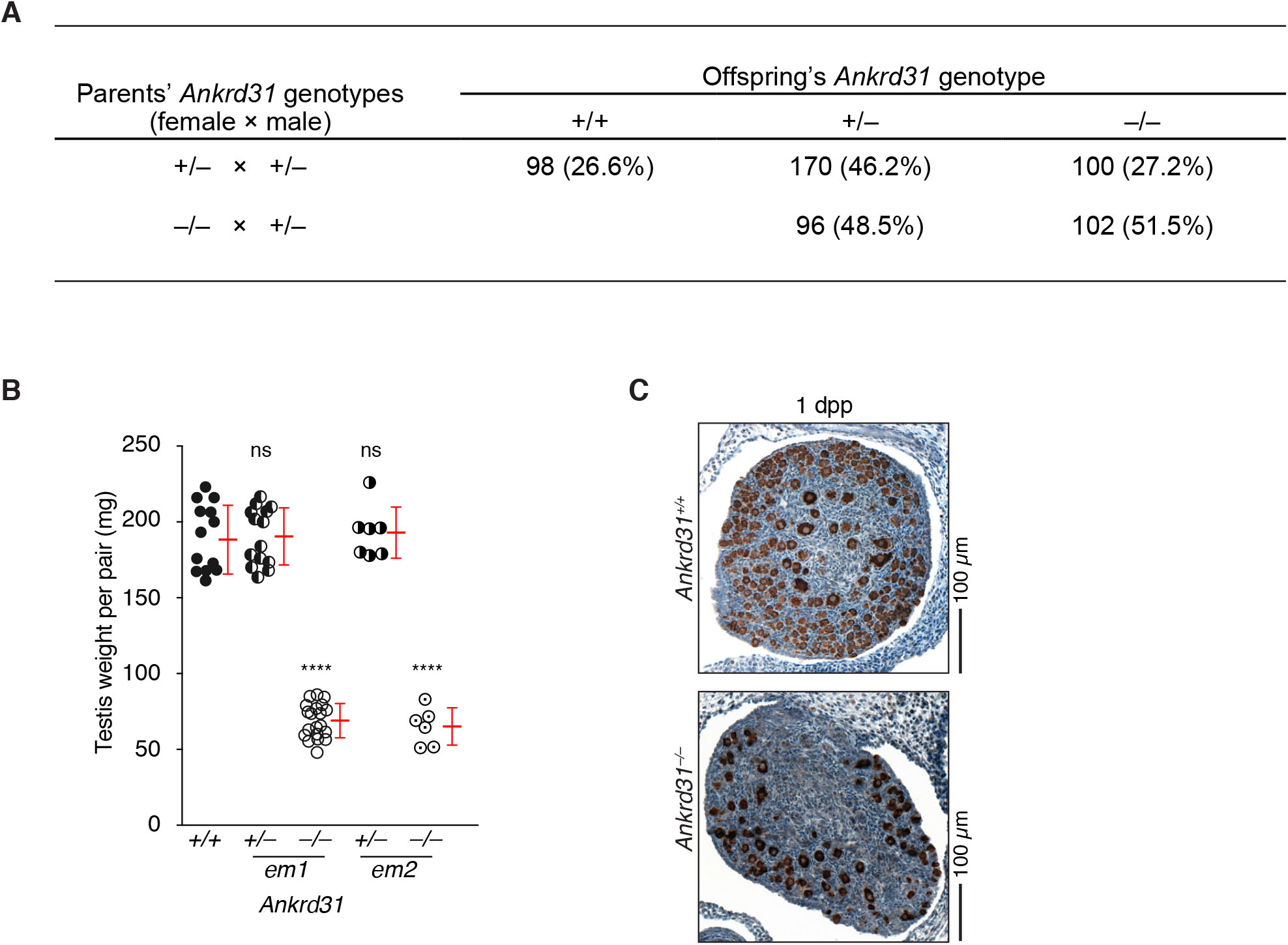
Compromised fertility of *Ankrd31^−/−^* mutants, related to Figure 2. (A) Mendelian segregation of the *Ankrd31* frameshift mutation. (B) Quantification of adult testis weights from mice heterozygous or homozygous for the *Ankrd31^em2^* allele. Data for wild type and mice heterozygous for the *Ankrd31^em1^* allele are reproduced from Figure 2A for comparison (red lines, mean ± s.d.). The results of two-tailed t tests for comparison to wild type are indicated: ns, not significant (p > 0.05); ****, p ≤ 0.0001. (C) Representative ovary sections from newborn animals immunostained for MVH to mark oocytes (brown stain).

**Figure S3.**
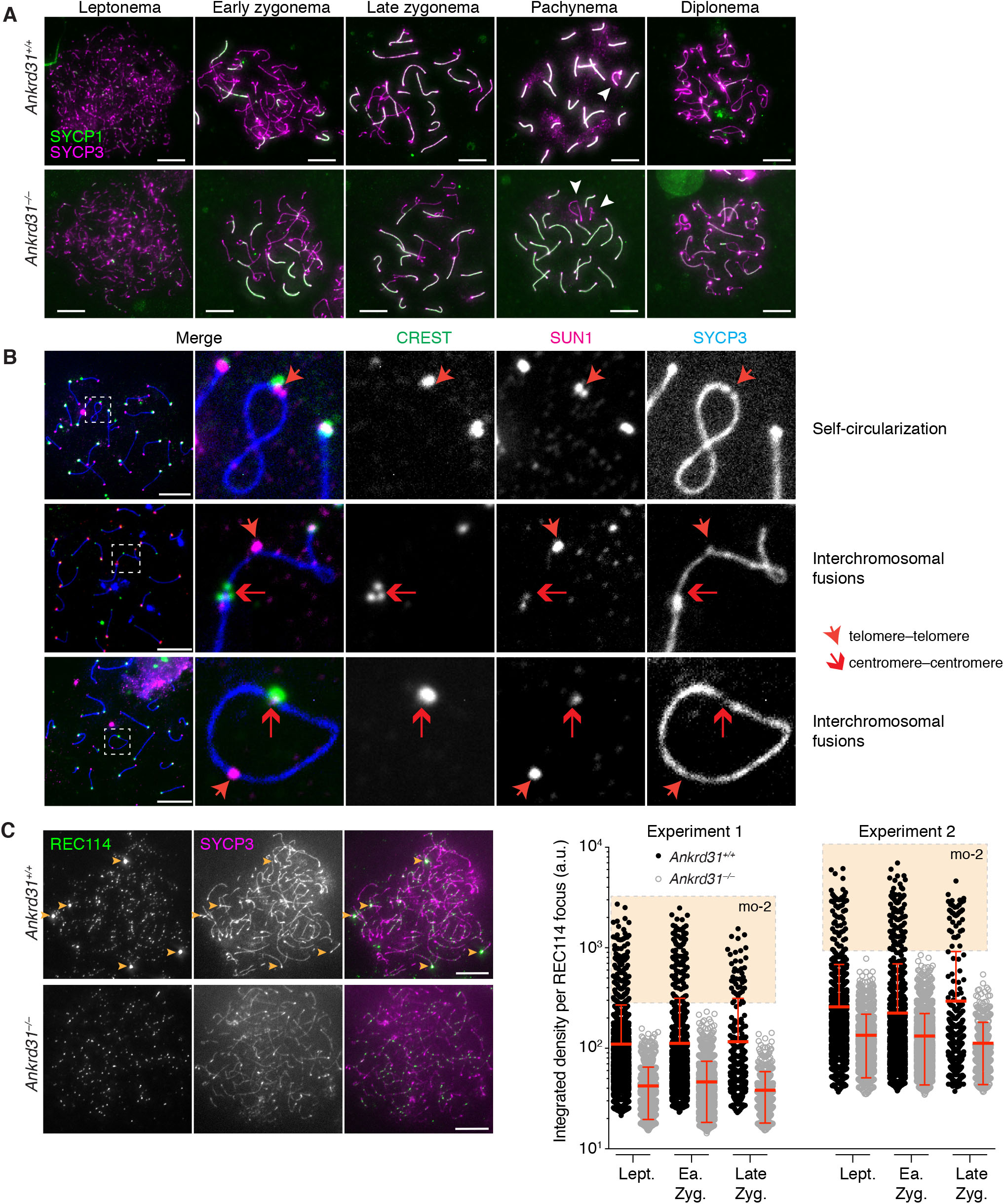
Synaptic defects and altered REC114 focus formation in *Ankrd31^−/−^* spermatocytes, related to Figure 3. (A) Meiotic progression in wild-type and mutant spermatocytes. Chromosome spreads were immunostained for SYCP1 and SYCP3. Arrowheads indicate sex chromosomes. (B) Additional examples of chromosome abnormalities observed at elevated frequency in pachytene spermatocytes from *Ankrd31^−/−^* mutants. (C) REC114 focus intensity in *Ankrd31^−/−^* spermatocytes. At left, additional views are provided of the same cells shown in Figure 3D. Arrowheads indicate REC114 blobs on mo-2-containing regions, including the PAR. At right are plots of integrated REC114 focus intensities for two independent experiments. The tan-shaded regions indicate where the heavily stained mo-2-associated blobs fall in the plots. The graph in Figure 3D summarizes the data from Experiment 1. Red lines indicate mean ± s.d. (downward error bars are not displayed for wild-type samples because they would be offscale on the log axis). Scale bars in all panels represent 10 μm.

**Figure S4.**
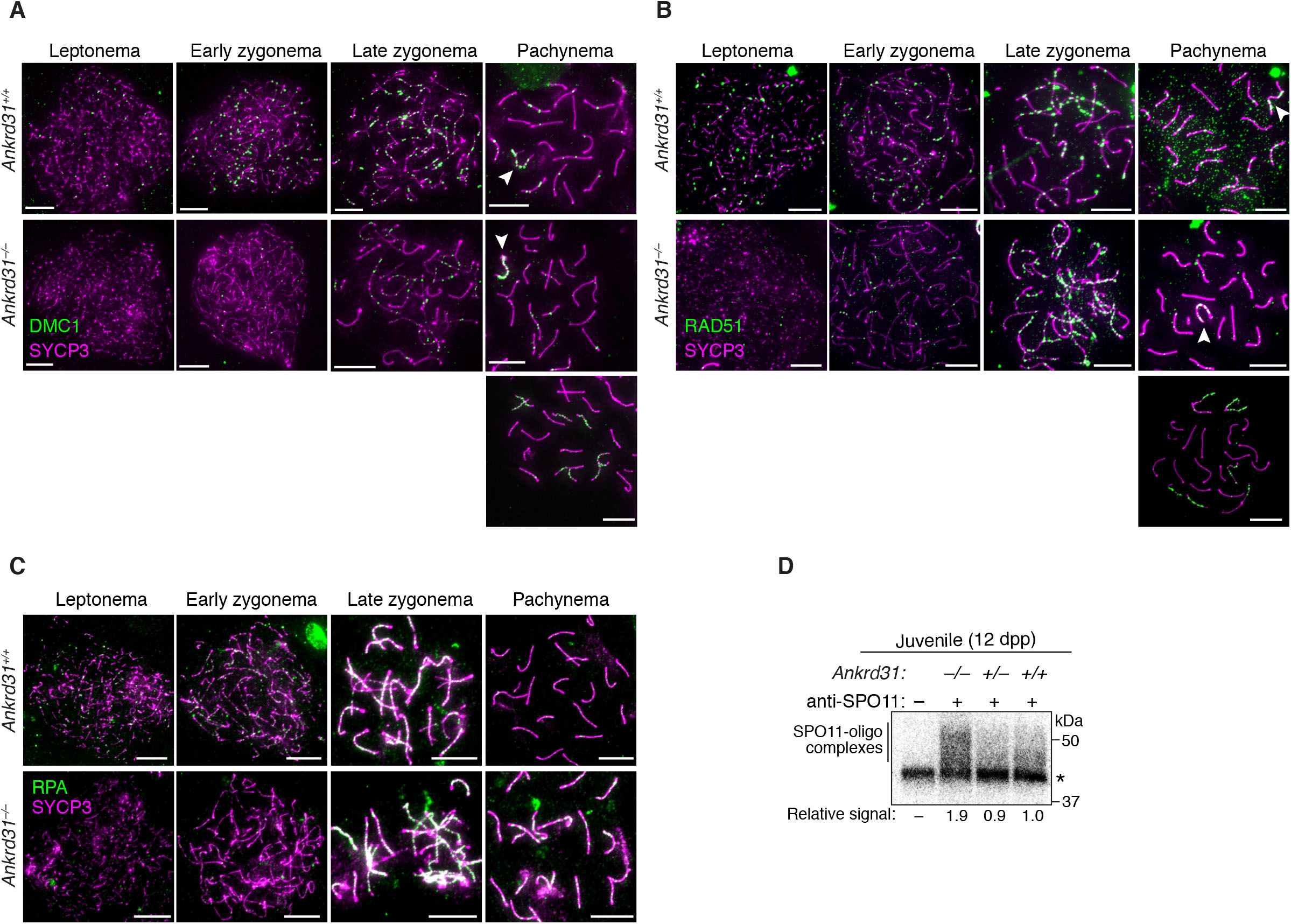
Dysregulated recombination in the absence of ANKRD31, related to Figures 3 and 4. (A) Altered numbers and timing of recombination foci. Additional images of DMC1 staining at different stages are shown. The early zygotene cells are reproduced from Figure 3E to facilitate comparison. Arrowheads on pachytene images indicate sex chromosomes. For the *Ankrd31^−/−^* mutant, a pachytene cell with normal-looking synapsis is shown, along with an example of a cell with autosomal asynapsis. (B) RAD51 immunostaining, as in panel A. (C) RPA immunostaining using antibodies to the RPA2 subunit, as in panel A. (D) Representative autoradiograph of SPO11-oligo complexes immunoprecipitated from juvenile testis extracts, radiolabeled with terminal transfer-ase and [α^32^P] dCTP, and separated by SDS-PAGE. Asterisk indicates nonspecific labeling. Quantified levels of radioactive signals were background-corrected and normalized to wild type.

**Figure S5.**
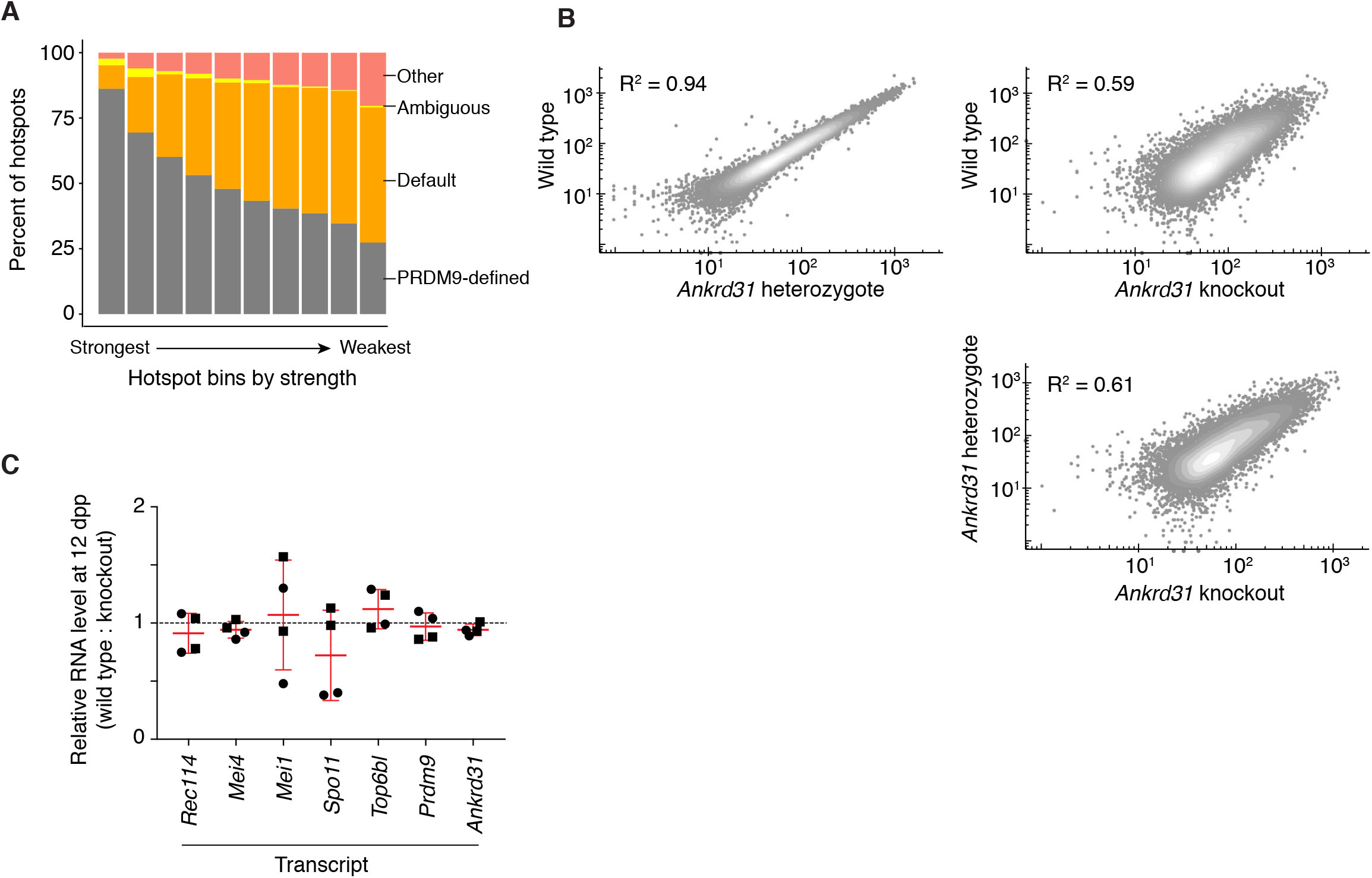
DSB distributions in *Ankrd31^−/−^* males, related to Figure 5. (A) Overlaps with PRDM9-defined and default hotspots. Hotspots called in the *Ankrd31^−/−^* juvenile SSDS map were split into ten bins by strength and scored for overlap with hotspots called from other datasets. “Ambiguous” sites are a hotspot in both B6 wild-type mice and in *Prdm9^−/−^* mice. “Other” sites are found in neither. (B) SSDS strength comparisons among strains at PRDM9-defined DSB hotspots (density gradient; grey (low) to white (high)). (C) *Prdm9* transcript levels are unaltered in *Ankrd31^−/−^* mutants. RT-qPCR was carried out on RNA extracted from wild-type or mutant animals at 12 dpp. Two primer pairs were used for each gene (filled circles and squares, respectively), for two independent experiments. Red error bars indicate pooled means ± s.d. Interestingly, *Ankrd31* transcript levels were also not reduced, suggesting that the mutant mRNA is not subject to substantial levels of nonsense-mediated decay, at least at this stage in development. *Spo11* transcripts were lower in two samples but not the others, so alterations in SPO11 expression are unlikely to be the cause of *Ankrd31^−/−^* phenotypes.

**Figure S6.**
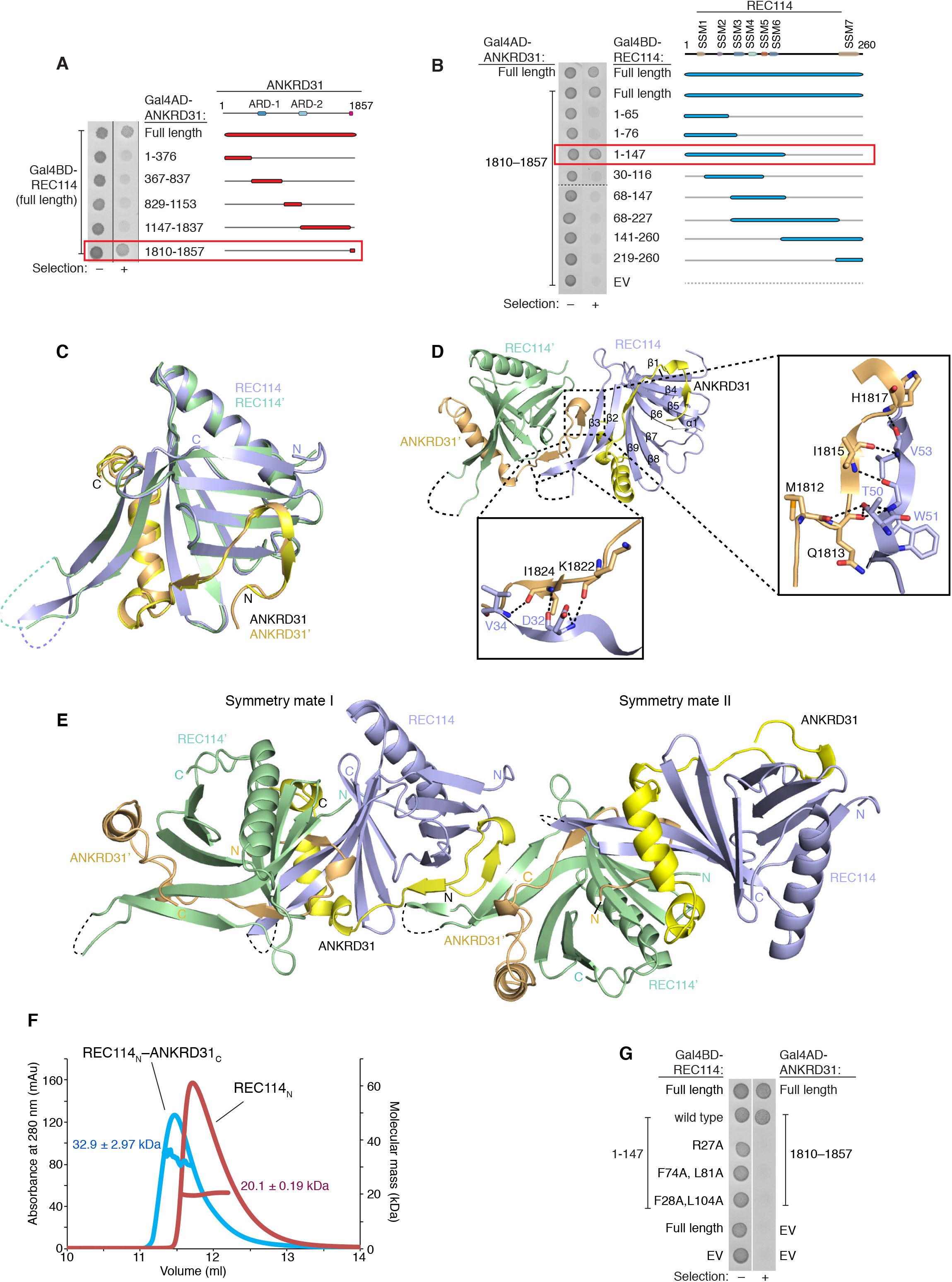
REC114_N_–ANKRD31_C_ complexes, related to Figures 6 and 7. (A) Identification of minimal interacting domain in ANKRD31. The indicated deletion constructs of ANKRD31 fused to Gal4 activating domain (AD) were assayed by yeast two-hybrid for interaction with full-length REC114 fused to Gal4 DNA binding domain (BD). Images show cell growth without (–) and with (+) aureobasidin selection for expression of the two-hybrid reporter. The C-terminal 48 amino acids of ANKRD31 is necessary and sufficient for interaction with REC114 (red box). (B) Identification of minimal interacting domain in REC114. Similar to panel A, the indicated deletion constructs of REC114 fused to Gal4-BD were assayed by yeast two-hybrid for interaction with ANKRD31 fused to Gal4-AD. The N-terminal 147 amino acids of REC114 is necessary and sufficient for interaction with ANKRD31 (red box). (C) Superposition of two REC114_N_–ANKRD31_C_ heterodimers in the asymmetric unit. (D) Detailed contacts across two REC114_N_–ANKRD31_C_ heterodimers in the asymmetric unit. (E) Interactions of two adjacent symmetry mate REC114_N_–ANKRD31_C_ dimers in the crystal lattice. (F) SEC-MALS analysis of REC114_N_ alone and of the REC114_N_–ANKRD31_C_ complex. (G) Yeast two-hybrid assay confirming interaction defects of the indicated mutant REC114 proteins.

**Figure S7.**
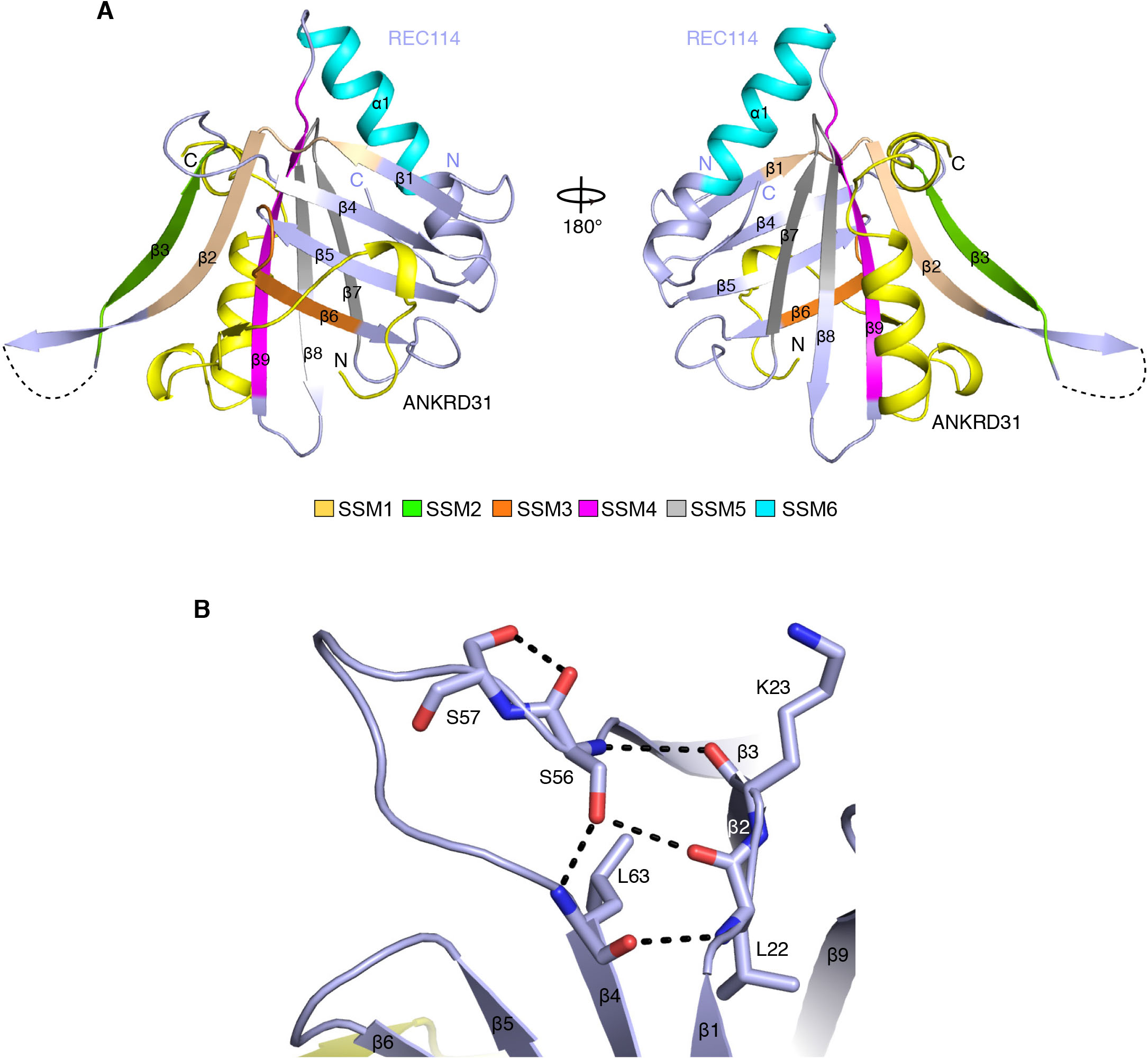
Conservation of REC114_N_ structural features, related to Figures 6 and 7. (A) Mapping of SSMs to the REC114_N_–ANKRD31_C_ heterodimer structure. Color code for SSMs is the same as in Figure 6B. (B) Detailed interactions involving S56 in REC114_N_.

**Table S1.** Data collection and structure refinement statistics, related to Figures 6, 7, S6, and S7.

|  |  |
| --- | --- |
| **Data collection** |  |
| Beamline | APS-24ID-C |
| Wavelength (Å) | 0.9791 |
| Space group | $P4_32_12$ |
| *Cell parameters* |  |
| *a, b, c* (Å) | 71.81, 71.81, 154.98 |
| $\alpha, \beta, \gamma$ (°) | 90, 90, 90 |
| Resolution (Å) | 50.0-2.80 (2.90-2.80)* |
| $R_{\text{merge}}$ (%) | 8.2 (77.5) |
| Average $I/\sigma(I)$ | 23.3 (1.9) |
| Completeness (%) | 99.9 (99.6) |
| CC (1/2) | 0.981 |
| Average redundancy | 7.5 (6.3) |
| No. of reflections | 10,273 |
| **Refinement** |  |
| $R_{\text{work}}/R_{\text{free}}$ (%) | 22.2/27.4 |
| No. of non-H atoms | 2612 |
| Average B factor (Å<sup>2</sup>) |  |
| REC114<sub>N</sub> | 84.8 |
| ANKRD31<sub>C</sub> | 90.5 |
| R.m.s. deviations |  |
| Bond length (Å) | 0.011 |
| Bond angles (°) | 1.157 |
| Ramachandran plot (%) |  |
| Favored | 95.1 |
| Allowed | 4.9 |
| Outliers | 0 |

\* Values in parentheses are for the highest resolution shell.

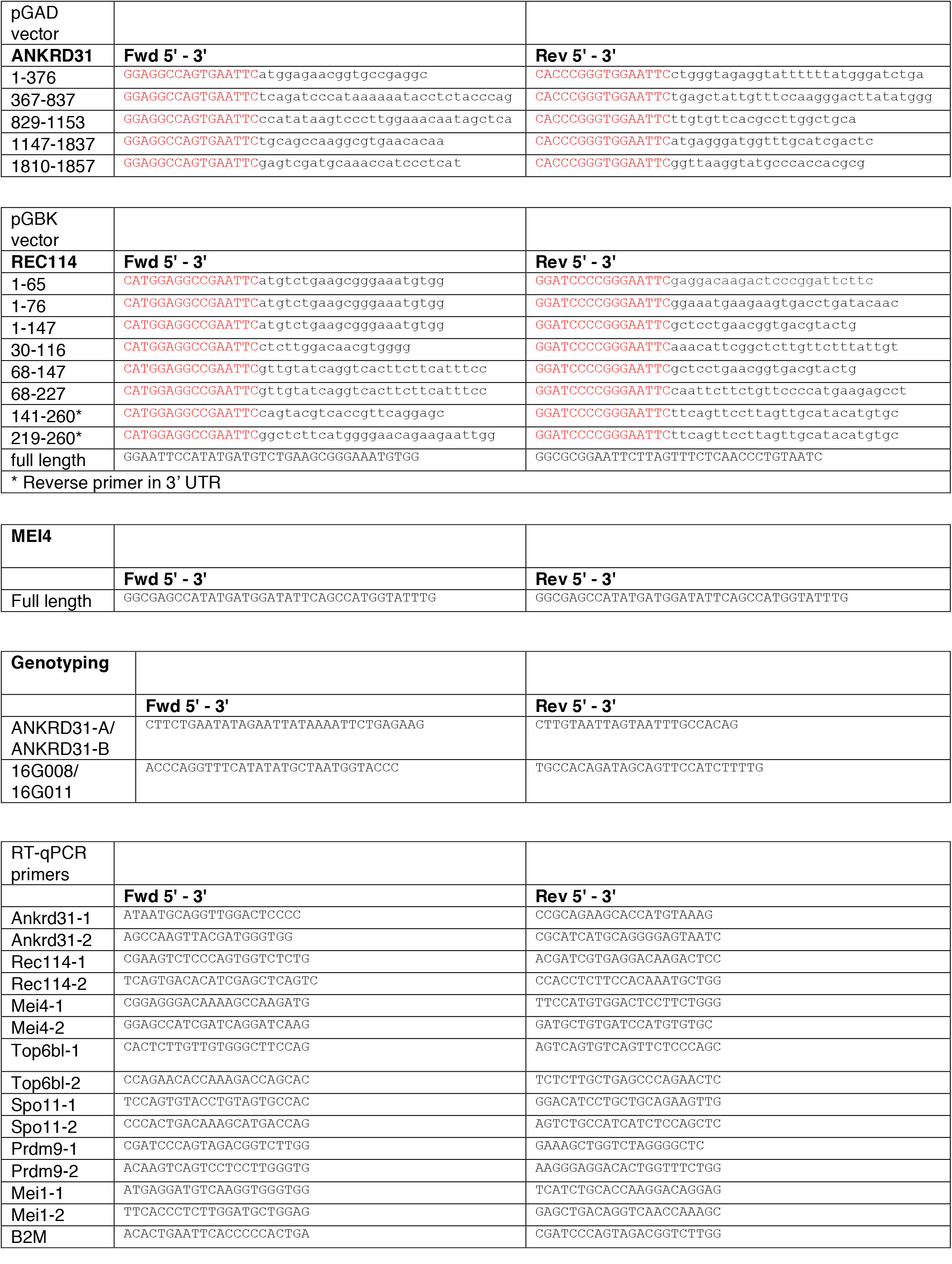

**Table S3.** Antibodies, related to STAR Methods.

| **Primary antibodies** |  |  |  |  |  |
| --- | --- | --- | --- | --- | --- |
| **Target** | **Dilution used for IF** | **Dilution used for Western** | **Host Species** | **Supplier** | **Catalog number** |
| γH2AX | 1:1000 | n/a | Rabbit | Santa Cruz | sc-101696 |
| DMC1 | 1:100 | n/a | Rabbit | Santa Cruz | sc-22768 |
| MLH1 | 1:25 | n/a | Mouse | BD-Pharmigen | 51-1327GR |
| RPA (H-100) | 1:75 | n/a | Rabbit | Santa Cruz | sc-28709 |
| SYCP3 (D-1) | 1:400 | n/a | Mouse | Santa Cruz | sc-74569 |
| SYCP3 (M-14) | 1:200 | n/a | Goat | Santa Cruz | sc-20845 |
| REC114 | 1:100 | n/a | Rabbit |  | n/a |
| ANKRD31 | 1:200 | 1:4000 | Guinea Pig | This study | n/a |
| ANKRD31 | 1:200 | n/a | Rabbit | This study | n/a |
| CREST | 1:500 | n/a | Human | Immunovision | HCT-0100 |
| SUN1 | 1:50 | n/a | Rabbit | Abcam | ab103021 |
|  | **Dilution for IP / Western** |  |  |  |  |
| SPO11-180 | 1:2000 |  | Mouse | MSKCC Antibody and Bioresource Core Facility | n/a |
|  | **Histology** |  |  |  |  |
| VASA/MVH/DDX4 | 0.17 microgram/ml |  | Rabbit | Abcam | ab13840 |
|  | **ChIP** |  |  |  |  |
| Histone H3K4me3 | n/a |  | Rabbit | Abcam | ab8580 |

| **Secondary antibodies** |  |  |  |  |  |
| --- | --- | --- | --- | --- | --- |
| **Target** | **Dilution used for IF** | **Fluorophore** | **Host Species** | **Supplier** | **Catalog number** |
| anti Goat | 1:400 | 350 | Donkey | Mol Probes | A-21081 |
| anti Guinea-pig | 1:400 | 488 | Goat | Mol Probes | A-11073 |
| anti Mouse | 1:400 | 594 | Donkey | Life Technologies | A21203 |
| anti Mouse | 1:400 | 594 | Goat | Mol Probes | A-11005 |
| anti Rabbit | 1:400 | 488 | Goat | Invitrogen | A11034 |
| anti Rabbit | 1:400 | 488 | Donkey | Thermofisher Scientific | R37118 |

